# Brazilin Removes Toxic alpha-Synuclein and Seeding Competent Assemblies from Parkinson Brain by Altering Conformational Equilibrium

**DOI:** 10.1101/2020.09.29.318220

**Authors:** George R. Nahass, Yuanzi Sun, Yong Xu, Mark Batchelor, Madeleine Reilly, Iryna Benilova, Niraja Kedia, Kevin Spehar, Frank Sobott, Richard B. Sessions, Byron Caughey, Sheena E. Radford, Parmjit Jat, John Collinge, Jan Bieschke

**Affiliations:** Colorado College, Colorado Springs, CO, USA; University College London Institute of Prion Diseases / MRC Prion Unit, London, UK; Washington University in St. Louis, St Louis, MO, USA; Astbury Centre for Structural Molecular Biology, School of Molecular and Cellular Biology, University of Leeds, Leeds, LS2 9JT UK; Biomolecular and Analytical Mass Spectrometry group, University of Antwerp, Antwerp, Belgium; Biomedical Sciences Building, University Walk, Bristol, BS8 1TD, UK; Rocky Mountain Laboratories, NIAID, NIH, Hamilton, MT, USA

**Keywords:** alpha-synuclein, Brazilin, Amyloid, Parkinson’s Disease, Polyphenol, Natural Compound, Neurodegeneration, RT-QuIC, Docking, Neurotoxicity

## Abstract

Alpha-synuclein (α-syn) fibrils, a major constituent of the neurotoxic Lewy Bodies in Parkinson’s disease, form via nucleation dependent polymerization and can replicate by a seeding mechanism. Brazilin, a small molecule derived from red cedarwood trees in Brazil, has been shown to inhibit the fibrillogenesis of amyloid-beta (Aβ) and α-syn, prompting our inquiry in its mechanism of action. Here we test the effects of Brazilin on both seeded and unseeded α-syn fibril formation and show that the natural polyphenol inhibits fibrillogenesis of α-syn by a unique mechanism that is distinct from other polyphenols and is also distinct from its effect on Aβ. Brazilin preserves the natively unfolded state of α-syn by stabilizing the compact conformation of the α-syn monomer over the aggregation-competent extended conformation. Molecular docking of Brazilin shows the molecule to interact both with unfolded α-syn monomers and with the cross-β sheet structure of α-syn fibrils. Brazilin eliminates seeding competence of α-syn assemblies from Parkinson’s disease patient brain tissue, and treatment of pre-formed fibril assemblies with Brazilin significantly reduces their toxicity in primary neurons. Our findings suggest that Brazilin has substantial potential as a neuroprotective and therapeutic agent for Parkinson’s Disease.

**Highlights:**

- The natural polyphenol Brazilin binds to monomeric, oligomeric and fibrillar α-syn
- Brazilin shifts the equilibrium away from aggregation-competent monomer conformations
- Brazilin inactivates seeding-competent α-syn isolated from Parkinson patients’ brains
- Brazilin detoxifies α-syn aggregation intermediates and stabilizes mature amyloid fibrils

**Graphical Abstract:** 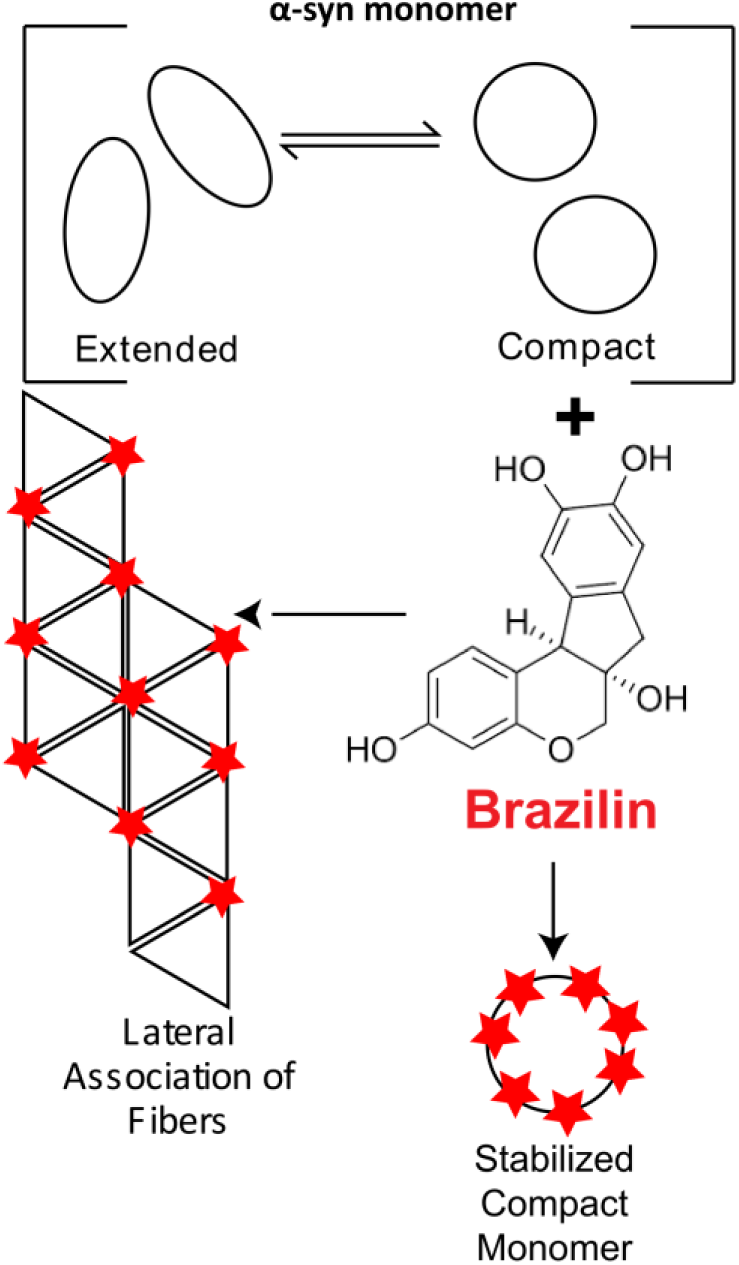

## INTRODUCTION

Parkinson’s Disease (PD), the second most common neurodegenerative disease following Alzheimer’s Disease (AD), is characterized by the successive loss of dopaminergic neurons in the substantia nigra pars compacta along with pathological aggregation of the natively unfolded α-synuclein (α-syn) protein into Lewy Bodies [1–3]. In its native state, α-syn is a 140 residue intrinsically disordered protein that can adopt an α-helical structure when bound to membranes either as part of the protein complexes or, possibly, through self-association [4,5]. Monomers of α-syn contain three regions with distinct compositions of amino acids (a.a.): the amphipathic N-terminal region (a.a. 1-60) which can adopt alpha helical structure, the hydrophobic non-amyloid-*β* component (NAC) region (a.a 61-95) responsible for the formation of β-sheet structure, and an unstructured C-terminal region (a.a. 95-140) which includes 14 acidic a.a. [6,7]

Formation of α-syn fibrils is a complex process with many intermediates including oligomers, ring-like oligomers, and protofibrils [6,8]. The fibrillization process for amyloids is a nucleation dependent pathway [9,10], where post nucleation species of oligomers and protofibrils act as seeds for monomers, which accelerate the formation of insoluble fibrils [4,11]. It has been suggested that endocytosis of the misfolded protein may be the initial step in the replication of the misfolded protein structures by prion-like mechanisms [12–14]. Toxicity of α-syn *in vivo* is related to the oligomeric forms of α-syn [15] as well as purified protofibrils and fibrils [16]. Processes that generate seeds will cause α-syn to autocatalytically replicate via secondary nucleation, a general mechanism, which is observed in other misfolded protein diseases such as AD and prion diseases [4,12]. Pathological α-syn assemblies can replicate *in vitro* and in tissues [17,18]. Mutant strains of α-syn, A30P, E46K, and A53T, have natively indistinguishable monomer conformations, but are associated with early onset PD, likely due to accelerated spherical oligomer and fibril formation [9,19,20].

In solution, α-syn exists in equilibrium between extended and more compact conformations. The dynamic ensemble between these two conformational families can be shifted by environmental factors and ligand binding, [21] and pH of solution along with temperature, crowding agents, membranes, metal ions and interacting proteins have also been shown to modulate the conformational states of α-syn [22]. It is well documented that polyvalent metals increase the aggregation potential of α-syn and lead to increased fibril formation of distinct morphology and toxicity [23–26], and a shift to the more compact form of α-syn upon metal binding has been observed by ESI-IMS-MS [27–29], suggesting a role for the more compact conformations of α-syn as a precursor to amyloid formation [30,31], while a transient α-helical monomer may be a precursor to an alternative nucleation pathway [32,33].

An extensive number of small molecules have been identified to inhibit amyloidogenesis in AD and PD (reviewed in: [34]). Recently, studies on Lacmoid, Orcein and its derivatives, Congo red and EGCG, have explored alternative therapeutic strategies, such as derailing amyloid formation, stabilizing mature fibrils and the depletion of oligomeric aggregation intermediates [4,21,35–39]. Hasegawa and coworkers investigated the effect of 79 compounds belonging to 12 chemical classes on the assembly of α-syn into fibrils, and identified seven classes (polyphenols, phenothiazines, polyene macrolides, porphyrins, rifamycins, Congo red and its derivatives, and terpenoids) containing strong fibril inhibitory effects [40]. Much effort has been specifically focused around the study of polyphenols. This class of compounds is an important beneficial constituent in the human diet found in a wide range of fruits, vegetables, and beverages including tea and red wine [41]. Polyphenols such as resveratrol, curcumin, and methylene blue have been implicated as treatment in many neurodegenerative disorders, as well as inflammatory and common ophthalmic disorders [42–44]. It is suggested that phenol rings have a different mode of stacking compared to benzene rings, and that this difference allows phenol rings to non-covalently bond the amyloidogenic aromatic residues of monomers / oligomers, as well inhibiting efficient fibril assembly [45]. It has also been proposed that the presence of vicinal hydroxyl groups (in the manner of three > two > one –OH) hinders the progress of the fibril self-assembly process and/or destabilizes its amyloid structure and that antioxidants within the natural polyphenols scavenge reactive oxygen species as well as inhibit deposition of α-syn fibrils in the brain [41]. The presence of vicinal hydroxyl groups is a common feature of some of the most effective natural inhibitors of α-syn aggregation. In an assay of 169 compounds screened to inhibit α-syn fibril formation, all but one of the inhibitors was a catechol containing compound, including dopamine, L-dopa, norepinephrine, and epinephrine [46].

Brazilin, a natural compound extracted from the wood of Caesalpinia sappan, was recently shown to have an inhibitory effect on the fibrillogenesis and cytotoxicity of Aβ [47] and hIAPP [48]. Aβ aggregation is the crucial event in the pathogenesis of AD, but soluble Aβ oligomers are believed to be the main neurotoxic agents [47]. Oligomeric aggregation intermediates of α-syn, which can form membrane pores, are likewise toxic in neuronal models [49,50]. The structure of Brazilin is similar to other natural compounds shown to have therapeutic effects on the aggregation of α-syn, namely the two vicinal hydroxyl groups and the two phenol rings (Figure 1a). The similarities of Aβ and α-syn aggregation combined with Brazilin’s structural similarities to other α-syn fibrillogenesis inhibitors as well as the inhibitory effects Brazilin has on Aβ, suggest that Brazilin could be a key candidate as a potential neuroprotective and therapeutic agent for PD. Very recently, it was confirmed that Brazilin inhibits α-syn fibril formation in vitro [51]. Here, we analyzed in detail the mechanism how Brazilin disrupts α-syn self-assembly and inactivates seeding competent α-syn assemblies derived from PD brain homogenates. Our biochemical and molecular findings suggest Brazilin to be a potential therapeutic and neuroprotective agent for PD with a novel mechanism of action compared to other polyphenolic compounds.

**Figure 1:**
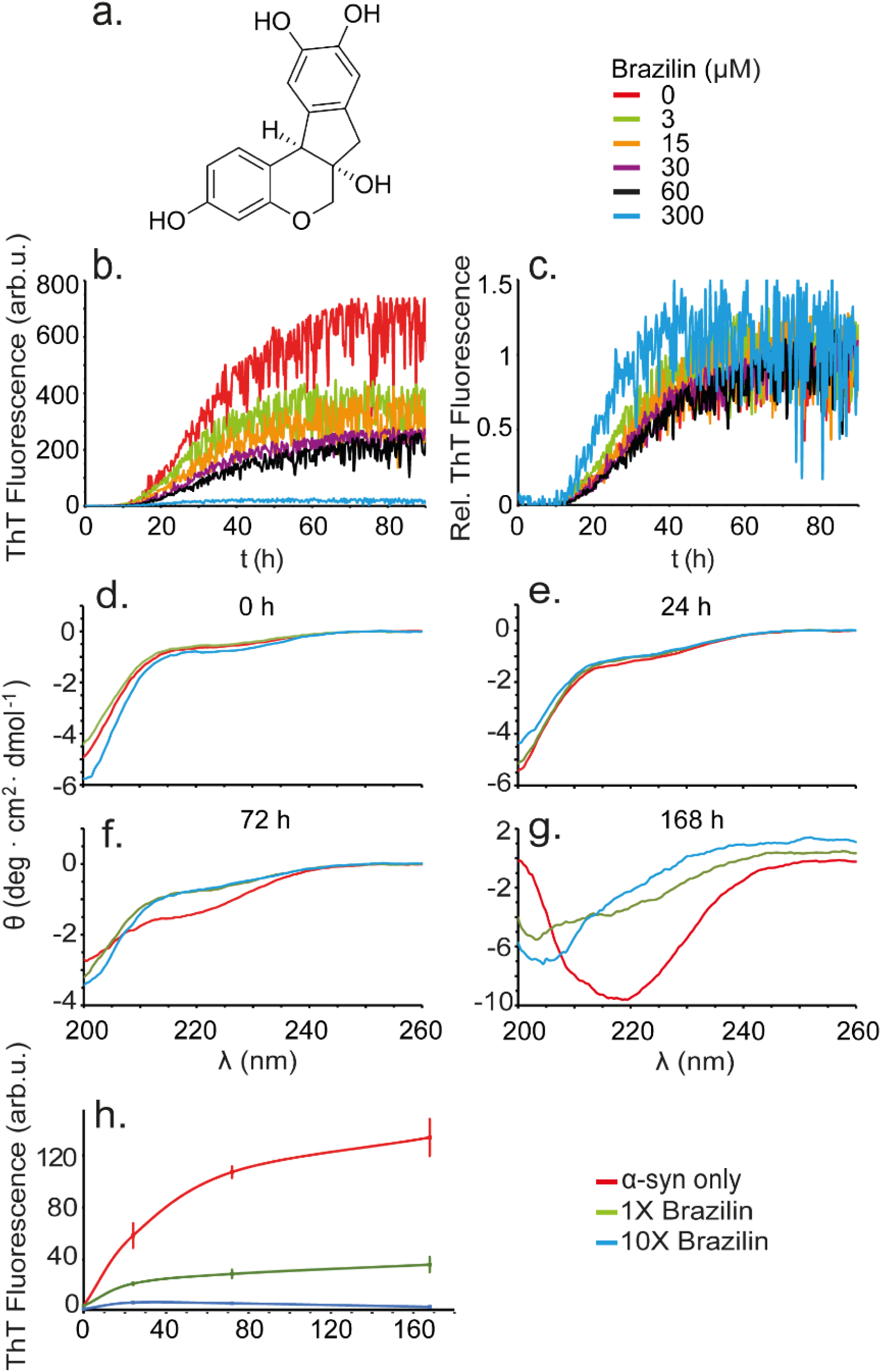
a) Structure of Brazilin molecule. b) Buffer subtracted ThT kinetic aggregation data of α-syn and Brazilin. c) ThT kinetic data normalized to the last ten hours of fluorescent readings. Graphs in (b) and (c) represent averages of triplicate curves with Brazilin concentrations from 0-300 μM as indicated by the color coding of graphs. d-f) Circular Dichroism (CD) spectra of 50 μM α-syn in ThT buffer with 0x (red), 1x (green), 10x (blue) molar ratio Brazilin at 0 h (d), 24 h (e), 72 h (f), and 168 h (g). h) ThT kinetics of CD samples after 2 minutes of sonication; means ± SD, n = 3.

## METHODS

### Alpha synuclein expression

α-Syn was expressed in E. coli as described previously [35]. Bacteria were grown at 37° C in LB and ampicillin (100 μg/mL). α-Syn expression was induced with IPTG once the OD 600 reached 1.1. The bacteria were then grown overnight at 30° C. The bacterial suspension was spun for 30 minutes at 3166 *×g* at 4°C. Two pellets of varying size were frozen at −80°C for 2 hours and then resuspended in 300 and 150 μL (large and small pellet respectively) of lysis buffer (10mM Tris HCl, 1mM EDTA, pH 8) containing Roche complete protease inhibitor. The solutions from the small and large pellets were then sonicated for 30 and 45 minutes respectively, boiled for 20 minutes, and spun down at 3185 *×g* for 30 minutes at 4°C. The resulting α-syn pellet was collected and dissolved in 25 mM Tris, pH 7.7. The α-syn was purified using a FPLC Resource Q anion exchange column with a salt gradient from 0–1 M NaCl. The protein was dialyzed overnight in ammonium bicarbonate buffer (10 mM, pH 7.5), spun down at 3185 *×g* for 30 minutes at 4°C, frozen and then lyophilized for storage.

Mutant K23Q α-syn for RT-QuIC assays was expressed in E. Coli and purified using an osmotic shock protocol as previously reported by Groveman et. al. (2018) [52].

### ThT aggregation assays

α-Syn (1 mg/mL) was dissolved in 10 mM NaOH and sonicated (VWR® Symphony^™^ Ultrasonic Cleaner) for 20 minutes. The α-syn was then spun down at 90,900 *×g* for 20 minutes at 4°C. The concentration of monomerized α-syn was then determined by OD_280_ using an extinction coefficient of 5960/M/cm. The α-syn was then aggregated in a non-binding 96 well plate (Corning, #3651) at a concentration of 30 μM with intermittent shaking in aggregation buffer (100 mM NaP, pH 7.4, 10 mM NaCl, 0.1% NaN_3_, and 20 μM ThT). A 2 mm diameter glass bead was added to each well to accelerate the aggregation through stirring. The plate was kept at 37° C and agitated by orbital shaking once every 1 minute for 5 seconds. The ThT fluorescence was recorded with an excitation wavelength of 436 nm and an emission wavelength of 482 nm in a fluorescence plate reader (Infinite F200, Tecan).

### Formation of α-syn Seeds for Seeded Aggregation Assays

α-syn seeds were generated by sonication of α-syn aggregates at different aggregation time points for 10 minutes in a water bath sonicator (VWR® Symphony^™^ Ultrasonic Cleaner). 5% (m/m) α-syn seeds were then added to fresh α-syn monomers and aggregated with ThT as described above.

### Atomic Force Microscopy

Aliquots of aggregated peptides (10 μL) were placed on clean, freshly cleaved grade V-1 mica (Cat#: 01792-AB, Structure Probe, Inc., USA). After 10 minutes, the solvent was wicked off by filter paper and the mica was washed 4 times with 20 μL of ultrapure (Milli-Q) water to remove salts and buffer from the sample. Samples were dried overnight, and AFM images were acquired in tapping mode on a Veeco Dimension 3100 machine (Bruker) with Bruker FESP tips. Images were visualized using the Bruker Nanoscope software v1.5.

### Ultracentifugation / Gel Electrophoresis

The α-syn samples (50 μM) incubated under constant shaking at 37°C taken at various time points (24, 72, 168 h) were centrifuged (TL-100, Beckman) at 50k RPM (100,000 *×g*) for 30 minutes. The initial sample, supernatant after ultracentrifugation, and pellet dissolved in the original sample volume of 1:1 PBS: SDS loading buffer were then analyzed by SDS-PAGE (Bio-Rad) at 160 V for 30 minutes. The pellets were washed with 1x PBS to eliminate residual supernatant. The gel was then washed with water, fixed for 20 min (50% EtOH, 20% acetic acid), washed again with water and stained with EZBlue staining reagent (Sigma-Aldrich) overnight. Additional water washes were performed to de-stain the gel.

### Circular Dichroism

The α-syn samples (50 μM) incubated under constant shaking at 37°C taken at various time points (24, 72, 168 h) were diluted 2:1 in aggregation buffer (100 mM NaP, pH 7.4, 10 mM NaCl, 0.1% NaN_3_, and 20 μM ThT). The 168 h incubation samples were sonicated for 2 minutes in a water bath sonicator (VWR® Symphony^™^ Ultrasonic Cleaner). Circular dichroism (CD) spectra were recorded between 200 and 260 nm with a step size of 1 nm in a CD spectrometer (J-720, Jasco, Japan)[4]. The CD spectra of solutions without α-syn were subtracted as background from the CD signals with α-syn to isolate the α-syn-specific changes. All spectra were the average of 5 consecutive scans for each sample.

### Native Electrospray Ionization-Ion Mobility-Mass Spectrometry

Protein samples with a final concentration of 20 μM for native electrospray ionization-ion mobility-mass spectrometry (ESI-IM-MS) experiments were prepared in 50 mM ammonium acetate buffer (pH 6.8). Small molecule stocks in 100% DMSO were diluted into the buffer solution to achieve the required final concentrations (molar ratio of α-syn and small molecule: 1:1, 1:2.5 and 1:5). The percentage of DMSO in the solution was 2% (v/v) for all of the samples.

ESI-IM-MS analysis was performed on a Synapt G1 HDMS instrument (Waters Corp., Wilmslow, UK) with travelling-wave ion mobility (TWIMS). All the samples were analysed using positive ionization ESI with a spray capillary voltage of 1.8 kV using home-made borosilicate glass capillaries with filaments (Puller: the Flaming/Brown Micropipette Puller; Coater: Emitech Sputter Coater). The following instrumental parameters were used: sampling cone 50 V; source temperature 80 °C; backing pressure 3.0 mbar; trap collision energy 5 V; extraction cone 1 V; trap DC bias 22 V, transfer collision energy 4 V. Data were acquired over the *m/z* range of 500-4000. Data were processed by using MassLynx V4.1 and Driftscope V2.5 software supplied with the mass spectrometer.

### Brain Homogenate Preparation

All subjects provided consent to clinical assessment under UCSD IRB-approved protocol #080012, and had consented to their brains being obtained at autopsy. Clinical assessment and autopsy brain analysis were performed as outlined in [52]. Brain homogenates (BH; 10% *w*/*v*) were prepared by homogenizing the tissue in PBS using a Bead Beater (Biospec Products; 11079110z) for 1 min at maximum speed. The homogenate was then spun at 2000 ×*g* for 2 min at room temperature and the supernatant was transferred to a new tube and stored at − 80 °C for α-syn RT-QuIC analysis. For α-Syn RT-QuIC testing, brain homogenates were serially diluted in PBS.

### α-Syn RT-QuIC Protocol

RT-QuIC reactions were performed in black 96-well plates with a clear bottom (Nalgene Nunc International). We preloaded plates with 6 glass or silica beads (1 mm in diameter, BioSpec Products or 0.8 mm, OPS Diagnostics, respectively) per well. For brain homogenate seeded reactions, 2 μL of indicated BH dilutions were added to wells containing 98 μL of the reaction mix to give final concentrations of 40 mM phosphate buffer (pH 8.0), 170 mM NaCl, 0.1 mg/mL (6 μM) of histidine-tagged α-syn (filtered through a 100 kD MWCO filter immediately prior to use), and 10 μM ThT. The plate was then sealed with a plate sealer film (Nalgene Nunc International) and incubated at 42 °C in a BMG FLUOstar Omega plate reader with cycles of 1 min shaking (400 rpm double orbital) and 1 min rest throughout the indicated incubation time. ThT fluorescence measurements (450 +/− 10 nm excitation and 480 +/− 10 nm emission; bottom read) were taken every 45 min.

### Primary Neuronal Culture and Toxicity Analysis

Primary neuronal cultures were derived from brains of inbred FVB/N mice. Hippocampi of E17 mouse brains were dissected in HBSS (ThermoFisher Scientific) supplemented with 1% L-glutamine, 1% HEPES and 1% pen-strep. Cells were dissociated using 0.25% trypsin+0.04% benzonase, triturated mechanically and counted using a Neubauer haemocytometer. Cells were plated in DMEM supplemented with 10% horse serum (#26050-88, Invitrogen) at 10K/well to the inner 60 wells of poly-L-lysine-coated 96 well plates (Greiner, 655936). At 1 h post-plating, DMEM medium was aspirated and exchanged for Neurobasal medium (21103049, ThermoFisher Scientific) supplemented with 0.25% Glutamax (35050061, ThermoFisher Scientific) 2% Gibco B27 supplement (17504044, ThermoFisher Scientific) and incubated at 37°C (20% O_2_, 5% CO_2_). FVB/N neurons were maintained in culture for 11-12 days prior to a 72 h treatment. Images of live cells were taken on an IncuCyte S3 reader (Sartorius) with a 20x objective in phase contrast. 4 views were captured in each well every 4 h.

Neurite lengths were evaluated using the NeuroTrack module of the IncuCyte S3 software package (rev 2019A) using the following parameters: cell body cluster segmentation 0.7; cleanup 0; cluster filter 0; neurite sensitivity 0.25; neurite width 1 μm. Detected neurite masks are highlighted in pink in images. Neurite length data were normalized to the initial (0 h) value for each well and means ± standard deviation were calculated from quadruplicate sample wells. ANOVA statistical analysis (two-tailed, equal variances) was performed in GraphPad Prism.

### Transmission Electron Microscopy

End-point samples from wt α-syn aggregation assays (5 μL) were loaded onto carbon-coated 300 mesh copper grids (Electron microscopy Sciences) that had been glow discharged for 40 seconds using an PELCO easiGLOW™ glow discharge unit (Ted Pella Inc., USA). Samples were left to bind for 1 min, blotted dry, washed in water (3 x 30 μL), blotted, and then stained with 10 μL Nano-W (methylamine tungstate) stain (Nanoprobes) for 1 min. Images were acquired on a FEI Tecnai T10 electron microscope (FEI, Eindhoven, NL). RT QuIC samples were collected for TEM imaging after 16 h of incubation. To collect solutions, a pipet tip was used to vigorously scrape the well surfaces and pipet the solution. 2-8 wells were pooled for each reaction condition and the solutions briefly sonicated. Ultrathin carbon on holey carbon support film grids (400 mesh, Ted Pella) were briefly glow-discharged before being immersed into droplets of the fibril solutions for 30–60 min at room temperature. Grids were sequentially washed three times in MilliQ water before being negatively stained with Nano-W stain and wicked dry. Grids were imaged at 80 kV with a Hitachi H-7800 transmission electron microscope and an XR-81 camera (Advanced Microscopy Techniques, Woburn, MA).

### Molecular Modelling

#### Setup of systems

1) Unfolded α-syn monomers were generated by selecting the ten best-energy structures from a brief minimization (7000 EMC steps) of a pool of 60000 randomly generated conformations using the *ab initio* peptide folding program RAFT[53] while weighting the residues psi/psi angle selection by secondary structure propensity. Repeating this process 100 times gave a population of 1000 viable random coil structures that were converted to all-atom models with SCWRL4 [54]. One hundred structures were chosen from the pool at random and each placed in a simulation box 2 nm larger than the protein in every dimension for MD. BUDE [55,56] was used in surface-scanning mode to locate the best ten non-overlapping Brazilin binding sites on an α-syn monomer as the start point for the 100 simulations of α-syn monomers bound to 10 Brazilin molecules. 2) A bespoke program was used to pack 64 of the monomer initial simulation boxes into the smallest cube giving a 46 nm cube. Performing this with and without Brazilin bound gave the starting points for the two large simulations to explore aggregation. 3) The fibril fragment containing 12 α-syn monomers was centered in a 15.67 nm cube. BUDE was used to define a 12 × 1.2 nm grid within the box and add Brazilin in random orientations to this grid. Brazilin molecules (222, not overlapping the protein) were selected at random.

#### Simulations

All molecular dynamics simulations were performed with GROMACS 2018 [57] under the following protocol: Hydrogen atoms were added to α-syn consistent with pH 7. The periodic-boundary boxes were filled with TIP3P waters and 0.15 M sodium chloride to neutrality. The system was parameterized with the amber99SB-ildn forcefield [58]. Brazilin was parameterized as a mixture of both tautomeric catechol mono-anions with the general Amber force field (GAFF) [59]. Short range electrostatic and van der Waals interactions were truncated at 1.4 nm and long-range electrostatics treated with the particle mesh Ewald method. An initial relaxation of 10000 steps of steepest descents energy minimization was performed. Subsequently, 0.2 ns of dynamics at 310 K was initialized while tethering the protein to its initial position, then the position restraints removed for production runs. The simulations were performed under periodic boundary conditions. The temperature was maintained at 310 K using the v-rescale thermostat and the pressure at 1 bar with the Berendsen barostat. Twin temperature baths were used, one for the protein and the other for the water and ions. Bond constraints were applied to the water (SETTLE) and the protein (LINCS) to allow a 2 fs timestep for the leap-frog integrator.

### Analysis

The simulation data were analyzed using the tools in the GROMACS suite and with bespoke programs written in-house.

## RESULTS

### Brazilin Does not Delay Spontaneous α-syn Aggregation but Inhibits Conformational Change of α-syn to *β*-Sheet

To examine the inhibitory effects of Brazilin on the aggregation of α-syn, aggregation was monitored using a thioflavin-T (ThT) fluorescence assay. ThT increases in fluorescence upon binding to β-sheet rich amyloid-like structures [36,60]. Inhibition efficiency was monitored by measuring the fluorescence of pure α-syn with respect to α-syn treated with various concentrations of Brazilin. Brazilin was added to monomeric α-syn (30 μM) in concentrations of 3, 15, 30, 60, and 300 μM and the sample allowed to aggregate for 90 hours. As shown in the non-normalized data (Figure 1b), Brazilin reduced the ThT fluorescence of α-syn in a concentration dependent manner. However, the presence of Brazilin did not influence the length of the lag phase of aggregation kinetics under these conditions, as can be seen when aggregation curves are normalized to the same amplitudes (Figure 1c).

These data could indicate that (a) Brazilin prevented α-syn incorporation into amyloid structures, (b) acted during the later stages of α-syn aggregation, permitting the formation of an early aggregate species with reduced ThT binding or (c) that it competitively inhibited ThT binding without affecting fibril formation [61]. In the latter case, it would be expected that α-syn aggregates would retain their β-sheet structure in the presence of the small molecule. We measured the secondary structure of α-syn in the presence and absence of Brazilin using circular dichroism (CD) over the course of one week to distinguish between these hypotheses. CD spectra of α-syn at t = 0 h and 24 h are consistent with a lack of secondary structure, as has been previously observed [4,35,49,62] (Figure 1d-e). Upon protein aggregation, CD spectra of untreated samples show a minimum at 218 nm while α-syn treated with Brazilin fully (10x) or partially (1x) retains the unstructured state (Figure 1f-g), which demonstrate that Brazilin inhibits the formation of β-sheet structures of α-syn *in vitro*. ThT fluorescence recorded from CD samples matches the inhibition of β-sheet formation (Figure 1h), excluding that ThT inhibition merely reflects competitive binding of the small molecule. ThT and CD data would be consistent with a mechanism, in which Brazilin suppresses a large percentage of amyloid formation at the primary nucleation stage, but does not substantially affect the lag phase, because overall kinetics are dominated by secondary nucleation processes.

### Brazilin Inhibits Fibril Formation of α-syn and Delays the Formation of Large Insoluble Aggregates

To test whether the reduction in ThT signal reflected a reduction in α-syn fibril formation, we imaged the samples analyzed in Figure 1d-h by atomic force microscopy (AFM; Figure 2a) and negative stain transmission electron microscopy (TEM; Figure 2b). After 24 and 72 h incubation, α-syn treated with 1x and 10x Brazilin show small amorphous aggregates, while untreated α-syn has begun to form full fibrils. At t = 168 hours, untreated α-syn has formed full fibrils (∼800 nm in length, ∼10 nm in height), while treatment with equimolar Brazilin reduced fibril size and only amorphous aggregates were detected for 10x molar ratio (Figure 2a). The incubation series was repeated, and transmission electron microscopy images were taken showing similar α-syn morphology over the time course of incubation (Figure 2b). These data show that fibril formation of α-syn is inhibited after a single treatment of Brazilin over a one-week incubation period.

**Figure 2:**
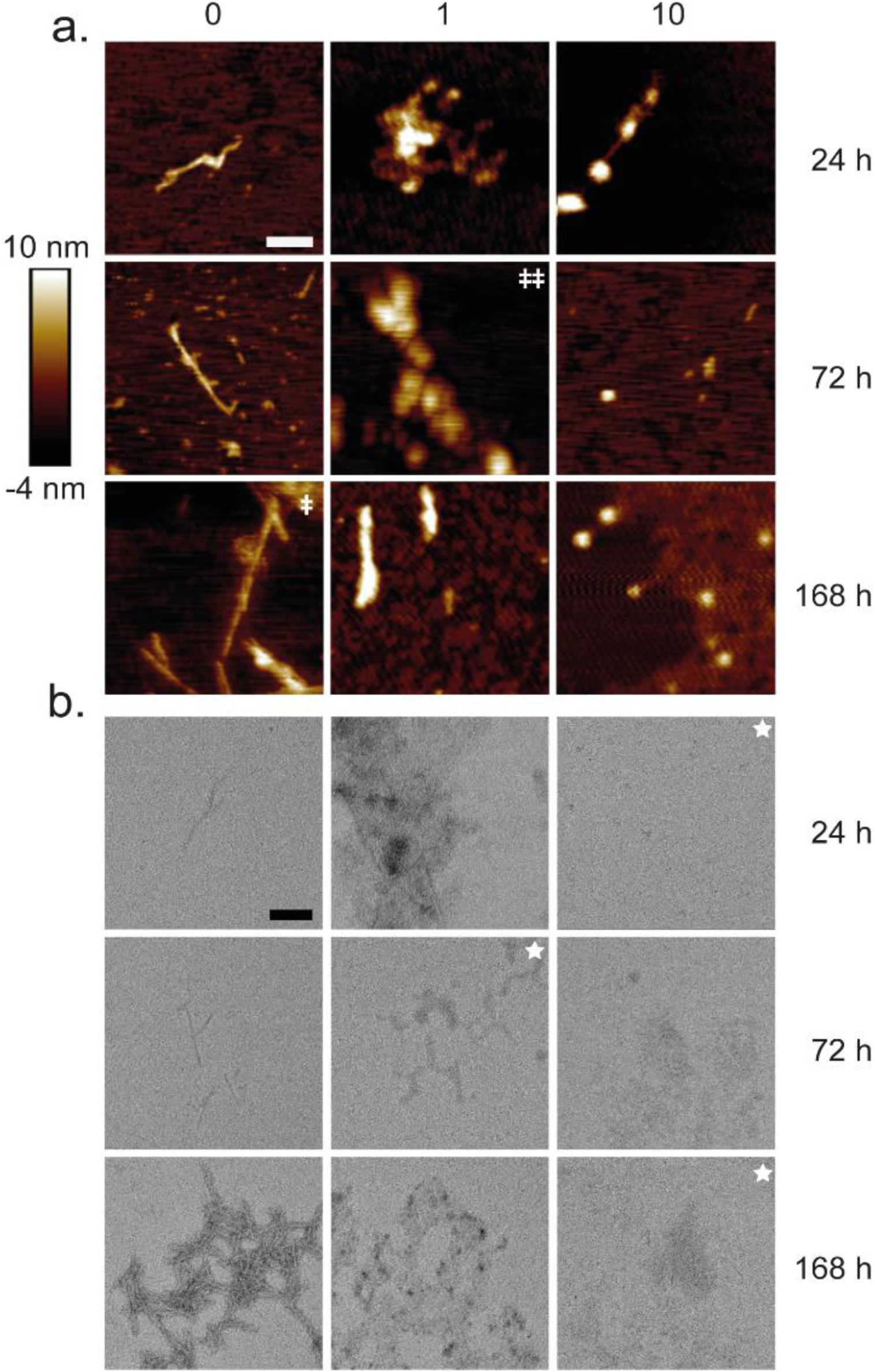
Atomic force microscopy (AFM) and transmission electron microscopy images taken at 24 h, 72 h, and 168 h time points of α-syn (50 μM) incubated with Brazilin (0x, 1x, 10x). All AFM heights are measured using the scale on the left unless otherwise noted within the panel (ǂ denotes height scale from −6 – 18 nm and ǂǂ denotes height scale from −4 −15 nm). The top left scale bar is 200 nm. Scale bar for TEM images is 200 nm, except for images containing a white star in the upper right corner, in which case it is 500 nm.

In order to quantify aggregation, the same samples used for CD and AFM were centrifuged at 100,000 *×g* and total (T), soluble (S) and pellet (P) fractions were analyzed b y SDS PAGE and Coomassie blue stain (Figure 3). Initially all α-syn was soluble and ran as a 14 kD monomer band. At t = 72 h, non-treated α-syn had almost completely aggregated, while α-syn treated with 1x and 10x Brazilin stayed soluble. Tenfold excess of Brazilin quantitatively prevented α-syn aggregation even after one-week incubation (Figure 3d). Taken together, the data from ThT fluorescence, CD, AFM, TEM and ultracentrifugation indicate that Brazilin prevents the formation of α-syn amyloid fibrils and keeps the protein in an unfolded state, which is most likely the monomer. Interestingly, a small amount of SDS-resistant α-syn dimer appeared in Brazilin treated samples, which may reflect the amorphous aggregates observed in AFM and TEM.

**Figure 3:**
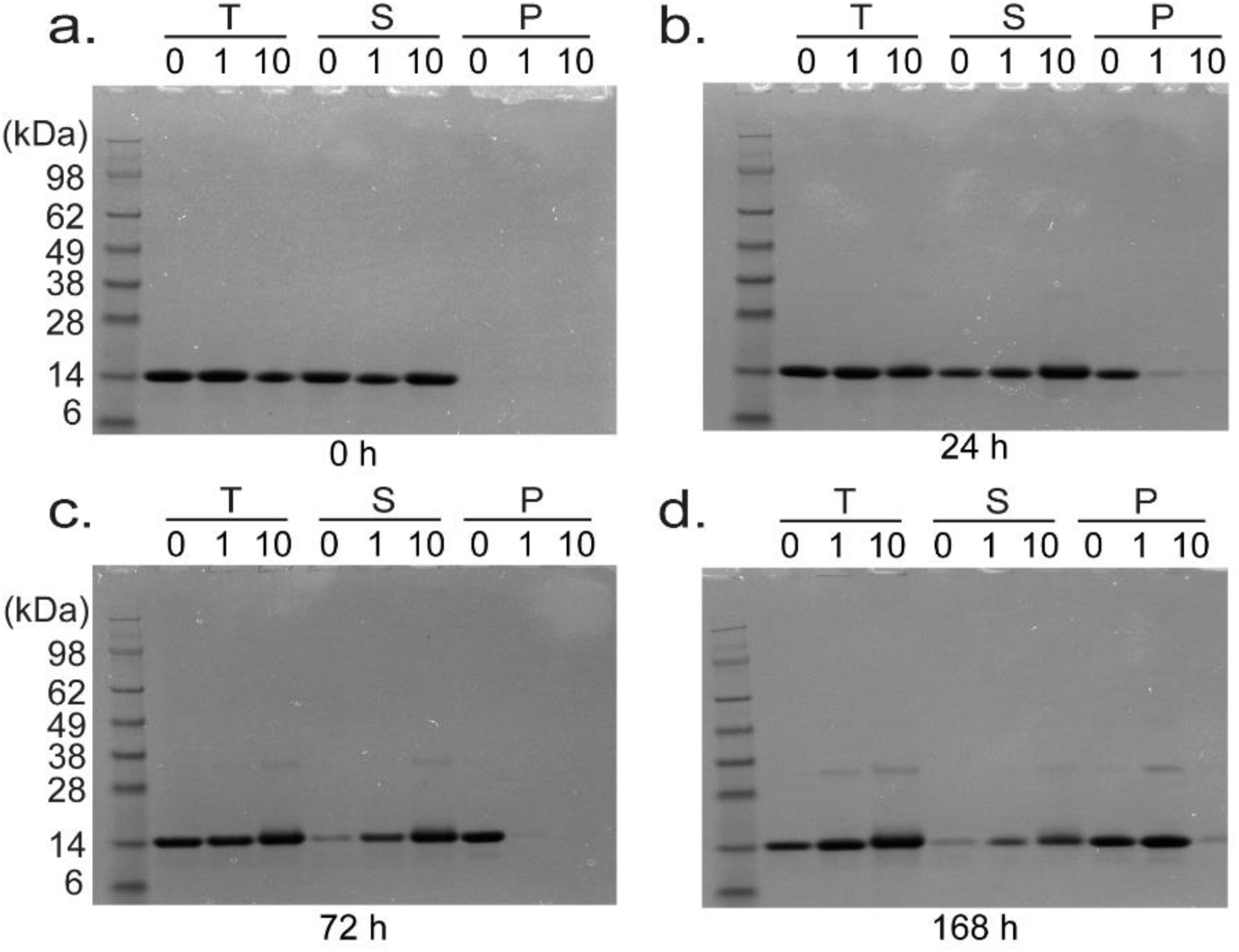
SDS Page gel electrophoresis of α-syn incubated with various concentrations of Brazilin at 0 h (a), 24 h (b), 72 h (c), and 168 h (d). The T represents total, S represents supernatant taken after ultracentrifugation at 100,000 *xg* at 4°C for 30 minutes, and P represents the pellet after ultracentrifugation after a wash with 1x PBS resuspended in an equal buffer volume.

### Brazilin Binds Preferentially to the Compact Conformation of α-syn

We analyzed α-syn by native ESI-IM-MS to determine how Brazilin prevented aggregation. The native mass spectrum of the sample treated with 2% DMSO only (same amount of DMSO in the Brazilin-treated sample) showed predominantly monomer species of α-syn (Figure 4a). Based on the bimodal charge state distribution observed in the spectrum, two major conformational families spanning from 15+ to 4+ were detected, reflecting extended and compact conformations of the protein, respectively (Figure S1) [21,30].

**Figure 4:**
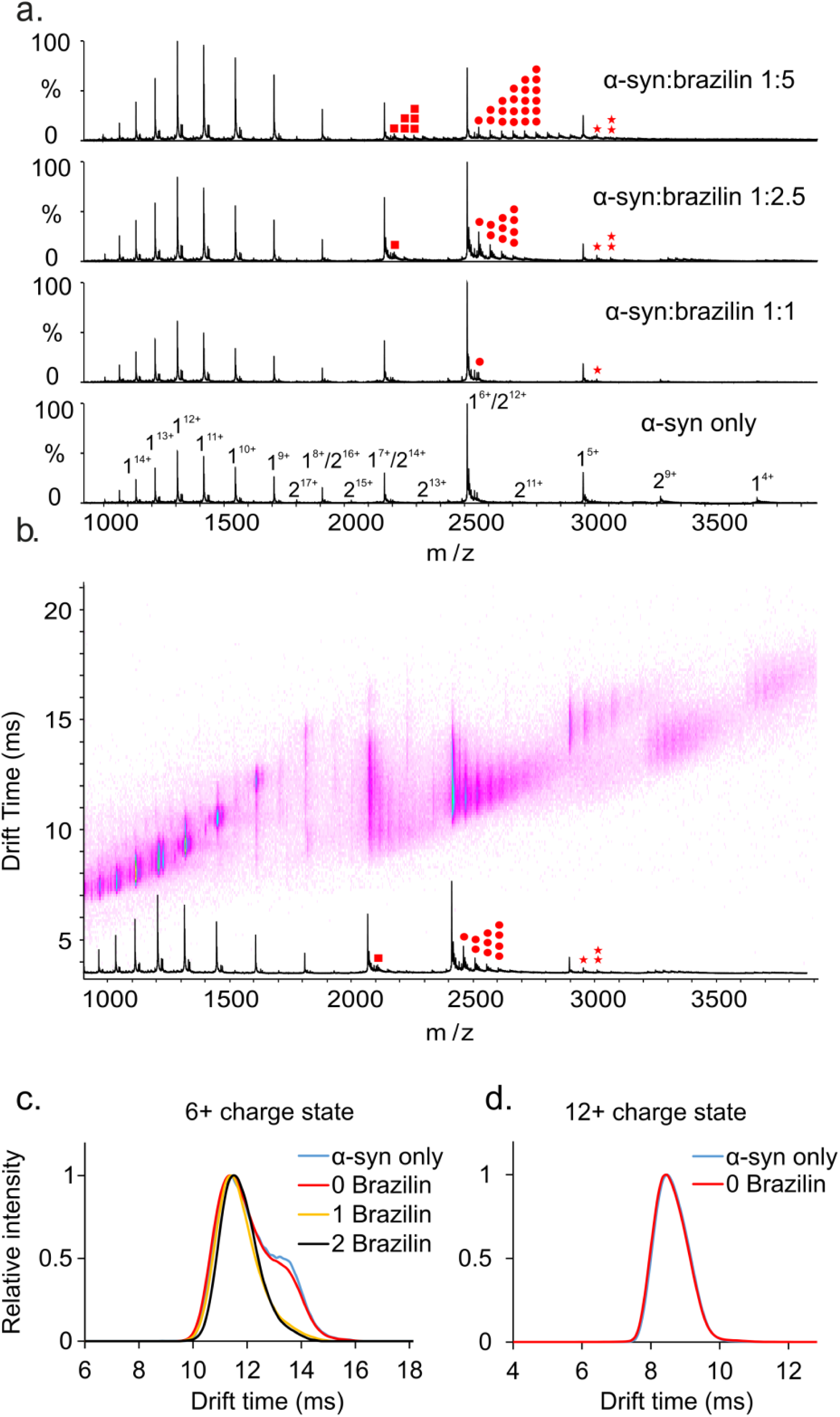
a) Positive-ion ESI mass spectra of α-syn in the absence (DMSO) or in the presence of 20 μM, 50 μM or 100 μM Brazilin in 50 mM ammonium acetate buffer. Brazilin binds to the 5+, 6+ and 7+ charge states of α-syn monomer. The monomer of α-syn is designated by “1”, the dimer by “2”. Brazilin copies are highlighted in red dots (7+: square, 6+: circle, 5+: star; the number of dots represents the number of ligands bound). b) Native ESI-IM-MS driftscope plot of different charge states of α-syn in the presence of 50 μM Brazilin (molar ratio of protein and small molecule is 1:2.5). The ESI-IM-MS driftscope shows IMS drift time versus *m/z*, and the corresponding ESI mass spectrum is shown at that bottom. Brazilin copies are highlighted in red dots as shown in Figure 4a. c) Arrival time distributions of the 6+ charge state of monomeric α-syn and its different Brazilin adducts as indicated in the inset (number of small molecules bound) obtained in the absence or presence of 50 μM Brazilin (α-syn:Brazilin molar ratio of 1:2.5. Blue line: monomer only; red line: monomer remaining ligand-free in the presence of Brazilin; yellow line: 1 Brazilin bound; black line: 2 Brazilin bound). Two distinct conformations of α-syn were observed for the unbound 6+ monomeric α-syn; while only one conformation was observed for the small molecule-bound ions. d) Arrival time distributions of the 12+ charge state of monomeric α-syn in the absence or presence of 50 μM Brazilin (α-syn:Brazilin molar ratio of 1:2.5. Blue line: monomer only; red line: monomer remaining ligand free in the presence of Brazilin).

As shown in Figure 4a, Brazilin bound preferentially to the low charge states of α-syn (5+ to 7+) with up to eight Brazilin ligands associated with a single molecule at the 1:5 (mol/mol) equivalent of α-syn and Brazilin. Notably, no brazilin bound to the more highly charged states representative of the extended form of the protein. Such a binding “selectivity” for the more compact lower charge states was also previously found for EGCG, whereas dopamine showed the exact opposite behavior by binding preferentially to the more extended highly charge states of α-syn [21]. It is possible that multiple molecules of Brazilin bind to each other forming a complex which then binds to the α-syn, but there were no complexes of free Brazilin observed in the mass spectra, implying that multiple molecules of Brazilin are interacting independently with α-syn monomers. To analyze the conformational specificity of Brazilin binding, the 6+ and 12+ charge states were chosen as representative of the high and low charge states of the protein, respectively, and their conformations were analysed by IM-MS. Drift time (dt) analysis revealed that there are two main conformations observed in the 6+ monomeric α-syn with drift times of 11.4 ms and 13.4 ms when the protein was treated with DMSO only (Figure 4c, α-syn only). When α-syn was incubated with Brazilin, the drift times of the 6+ charge state for the unbound α-syn remained unchanged, indicating that the small molecule does not induce a detectable conformational change of the protein (Figure 4c, 0 Brazilin). The same results were observed with all the charge states (the 12+ charge state is shown in Figure 4d). When analyzing the drift times of small molecule-bound ions (1 Brazilin bound and 2 Brazilin bound), the small molecule bound predominantly to the compact form of the 6+ charge state of α-syn with drift times of approximately 11.4 ms. Collectively, these data suggest that Brazilin preferentially binds to the low charge states of α-syn (i.e. 5+ to 7+), which is similar to the way in which EGCG binds to the protein. Multiple conformations were observed for the low charge states by IM-MS analysis. Specifically, two conformers were detected for the 6+ charge state and Brazilin binds predominantly to the compact conformation of the 6+ charge state of α-syn.

### Brazilin Does not Prevent the Recruitment of α-syn Monomers into Amyloid Fibrils

Preformed fibril seeds obviate nucleation [63], allowing us to analyze to effect of Brazilin on fibril growth. Solutions of α-syn (30 μM) were prepared with 5% (w/v) seed, formed by sonication of α-syn fibrils, and 3, 15, 30, 60, and 300 μM concentrations of Brazilin to test whether the small molecule prevented the recruitment of α-syn monomers to preformed fibrils (Figure 5a, b). Similar to *de novo* aggregation, Brazilin reduced the amplitude of ThT fluorescence in a concentration dependent manner. However, after normalizing ThT amplitudes to the final ThT amplitude, it was clear that Brazilin concentration has no effect on the seeded aggregation of α-syn. We calculated apparent growth rates from the initial slopes of ThT kinetics (Figure 5c, d). These growth rates were similar across all concentrations of small molecule when normalized for ThT amplitudes. We quantified total (T), soluble (S) and aggregated (P) α-syn from the end point of the seeding assay by ultracentrifugation and SDS-PAGE (Figure 5e). Their relative fractions remained unchanged by Brazilin, confirming that the small molecule did not directly inhibit the incorporation of α-syn monomer into growing fibrils.

**Figure 5:**
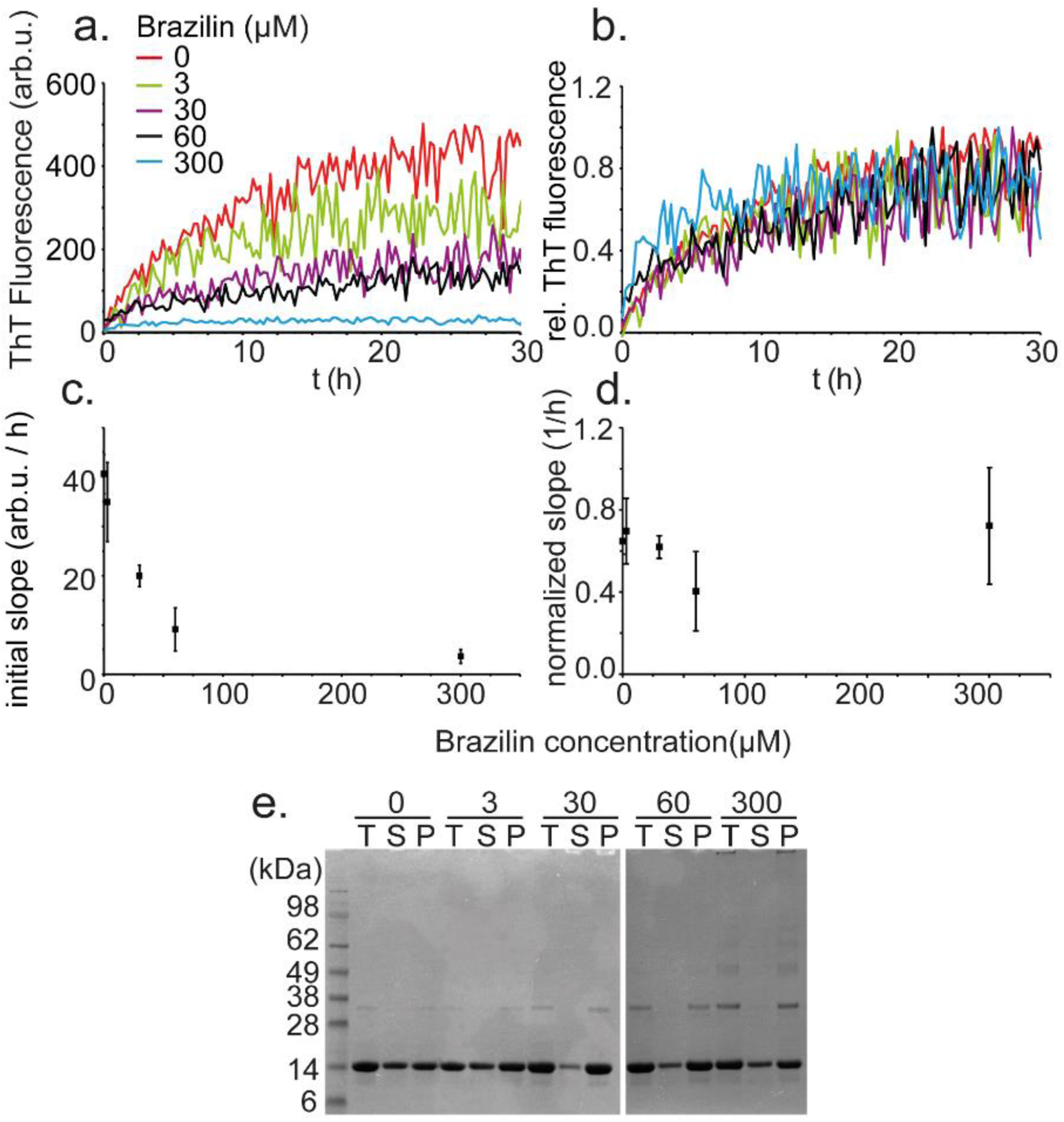
a) Buffer subtracted ThT kinetic aggregation data of α-syn (30μM) in ThT buffer with various concentrations of Brazilin in the presence of 5% (w/w) fibrillar seed. Seeds were generated by sonication of mature fibrils for 10 minutes. b) Buffer subtracted ThT aggregation data from (a) normalized to the last ten hours of fluorescent readings. c), d) Average slope of the ThT kinetics of a) and b), respectively. e) SDS gel electrophoresis of seeded α-syn aggregation in the presence of various Brazilin concentrations. The T represents total, S supernatant, and P represents the pellet fraction taken after ultracentrifugation of the total at 100,000 *xg* at 4°C for 30 minutes, dissolved in 1:1 NaP:SDS loading buffer after a wash with 1x PBS.

### Brazilin Inactivates Synthetic and Brain-derived α-syn Seeds

We then tested whether Brazilin could affect the activity of α-syn seeds themselves. α-Syn fibrils (30 μM monomer) were formed at 37 °C in phosphate buffer in a fluorescence plate reader and Brazilin was added after 65 h incubation (Figure 6a). We observed a gradual concentration dependent decrease in ThT fluorescence (t_½_ ∼ 3h) upon addition of Brazilin as compared to the control sample, which may indicate a structural remodeling of the fibrils.

**Figure 6:**
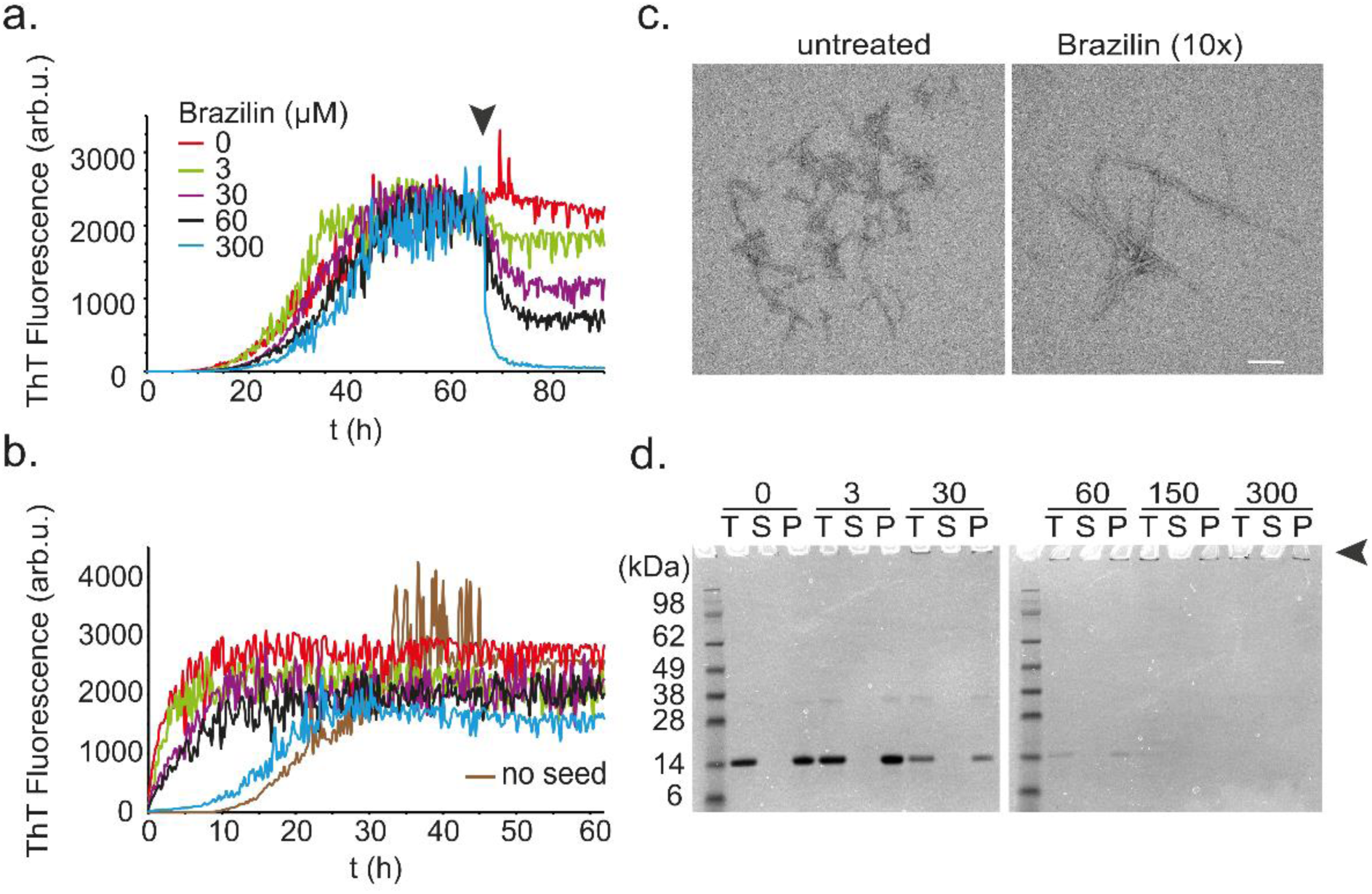
a) Buffer subtracted ThT kinetic aggregation data of α-syn (30μM) in ThT buffer with various concentrations of Brazilin added to mature fibrils. Graphs represent averages of triplicate curves. b) Buffer subtracted seeded aggregation of α-syn using various concentrations of Brazilin remodeled fibrils as seeds (5%) in ThT buffer. Seeds were created by sonication on ice in a water bath for 15 minutes. c) Negative stain TEM images of 0μM Brazilin and 300μM Brazilin remodeled fibrils. d) SDS Page gel electrophoresis of mature α-syn remodeled with various concentrations of Brazilin. The T represents total, S represents a fraction of the supernatant taken after ultracentrifugation of the total at 100,000 *xg* at 4°C for 30 minutes, and P represents the pellet left after ultracentrifugation dissolved in 1:1 NaP:Loading buffer after a wash with 1x PBS.

The end products of Brazilin treatment were then sonicated in a water bath for 15 minutes to generate seed, which were added to fresh α-syn monomer (Figure 6b). Treatment with Brazilin reduced seeding capacity in a concentration-dependent manner, indicated by the increase in lag phase when compared to untreated seeds, with 10x Brazilin having aggregation kinetics similar to that of untreated α-syn. In order to ensure that the effect was not due to residual Brazilin present in the seed solution from seed generation, we compared seeding kinetics of untreated fibrils in the presence and absence of Brazilin (15 μM, equivalent to the final concentration for seeds treated with 300 μM Brazilin), but found no effect on fibril growth rate, when normalizing for the reduction in ThT amplitude caused by the compound (Figure S2).

However, TEM revealed that amyloid fibrils of α-syn were still present after Brazilin treatment (Figure 6c) and quantification of aggregated α-syn via ultracentrifugation revealed no solubilization of α-syn fibrils by the Brazilin treatment, but rather the presence of fibril clusters (Supplementary Figure 3, Figure 6d). Rather, we found pelleted α-syn in insoluble high molecular weight aggregates that did not enter the SDS-gel after Brazilin treatment. This suggests that Brazilin, rather than dissolving amyloid fibrils, promoted the formation of large fibrillar assemblies, which were more stable and less seeding competent than untreated α-syn fibrils. Similar SDS-resistant high molecular weight aggregates were observed at high Brazilin concentrations in the previous seeding experiment (Figure 5e), supporting the interpretation that Brazilin coats and stabilizes large α-syn fibrils.

Finally, we tested the capability of Brazilin to inactivate α-syn from Parkinson’s disease brain homogenates. Seeds of α-syn can be amplified *in vitro* and detected via ThT fluorescence in real-time quaking-induced conversion assays (RT-QuIC; Figure 7a) [17,52]. Brain homogenates (10^−4^ dilution in PBS) from Parkinson patients (PD 14651, 4339, 3089) were added to monomeric α-syn K23Q (6 μM) with Brazilin (6 μM) and incubated for 48 h. Here, the seed concentration provided to the reaction can be estimated to be roughly 10^8^-fold lower than the seed concentration used above in Figure 5 based on comparisons of end-point dilution RT-QuIC titrations of Parkinson’s disease brain and known quantities of synthetic α-syn fibrils [52]. Under these conditions, Brazilin treatment prevented the induction of ThT positive aggregates. Control brain homogenates from control corticobasal degeneration (CBD) patients did not induce ThT fluorescence. Imaging by TEM confirmed that Brazilin treatment prevented the formation of fibrillar α-syn aggregates in RT-QuIC (Figure 7b). Conversely, RT-QuIC reactions seeded with CBD brain homogenates did not show α-syn fibrils (Figure S3). Brazilin was also tested in an RT-QuIC reaction using a dilution series of PD brain 3089 at a sub-stoichiometric concentration relative to α-syn monomers, and was found to completely inhibit the seeding of α-syn relative to positive controls, demonstrating potent inhibition of fibril formation (Figure S4a, b). The potent inhibition of seeding under RT-QuIC condition seemingly contradicts the observation that Brazilin does not directly inhibit fibril elongation (Figure 5). However, with the much lower initial seed concentration in the RT-QuIC reactions, we assume that fibril elongation alone, i.e., without the exponential seed amplification provided by secondary nucleation via lateral nucleation of fibril formation, would not be detectable. Thus, these data are consistent with the notion that Brazilin markedly reduced the activity of Parkinson’s-associated α-syn seeds by reducing secondary nucleation on fibril surfaces, even if, as suggested by data in Figure 5, primary fibril elongation was unimpeded.

**Figure 7:**
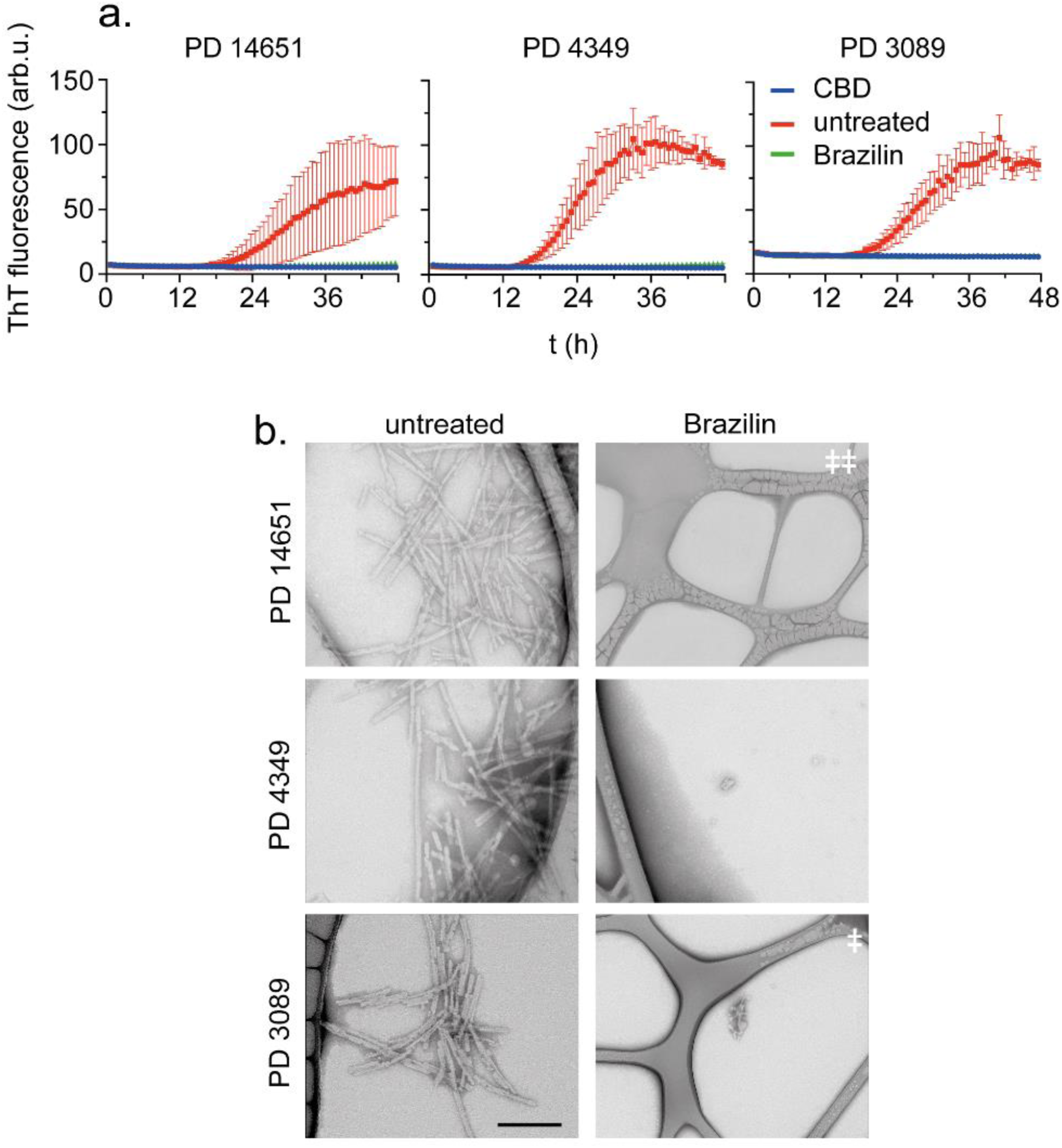
a) Inhibition of α-syn seeding activity by Brazilin in RT-QuIC using 2 μL of brain homogenate from three PD patient isolated post-mortem and 6 μM K23Q mutant α-syn as substrate in ThT Buffer. Graphs represent average of quadruplicate curves and each reaction was performed using brain dilution of 10^−4^. CBD represents brain homogenate from post mortem confirmed corticobasal degeneration cases b) Transmission electron microscopy images of RT-QuIC end products from various PD patient brains. Scale bar is 200 nm unless denoted otherwise within panel (ǂ = 500 nm, ǂǂ = 1 μm).

### Brazilin Removes α-Syn Toxicity in Primary Neurons

We then tested whether Brazilin could reduce the toxicity of preexisting α-syn assemblies using a novel method, which monitors neurite length of primary mouse hippocampal neurons in response to amyloid aggregates [64]. Fibrils were treated with Brazilin for 24 h at 1x, or 2x molar ratios (Figure 8a) and then added to neurons at 1 μM monomer equivalent concentration. Live neurons were imaged every 6 h and their neurite lengths were quantified relative to their pre-treatment state (Figure 8a, b). After 72 h incubation, α-syn treatment significantly reduced neurite lengths, whereas neurite lengths with Brazilin treated fibrils were not significantly different from untreated controls. Brazilin itself and monomeric α-syn did not reduce neurite lengths compared to buffer controls (Figure S5).

**Figure 8:**
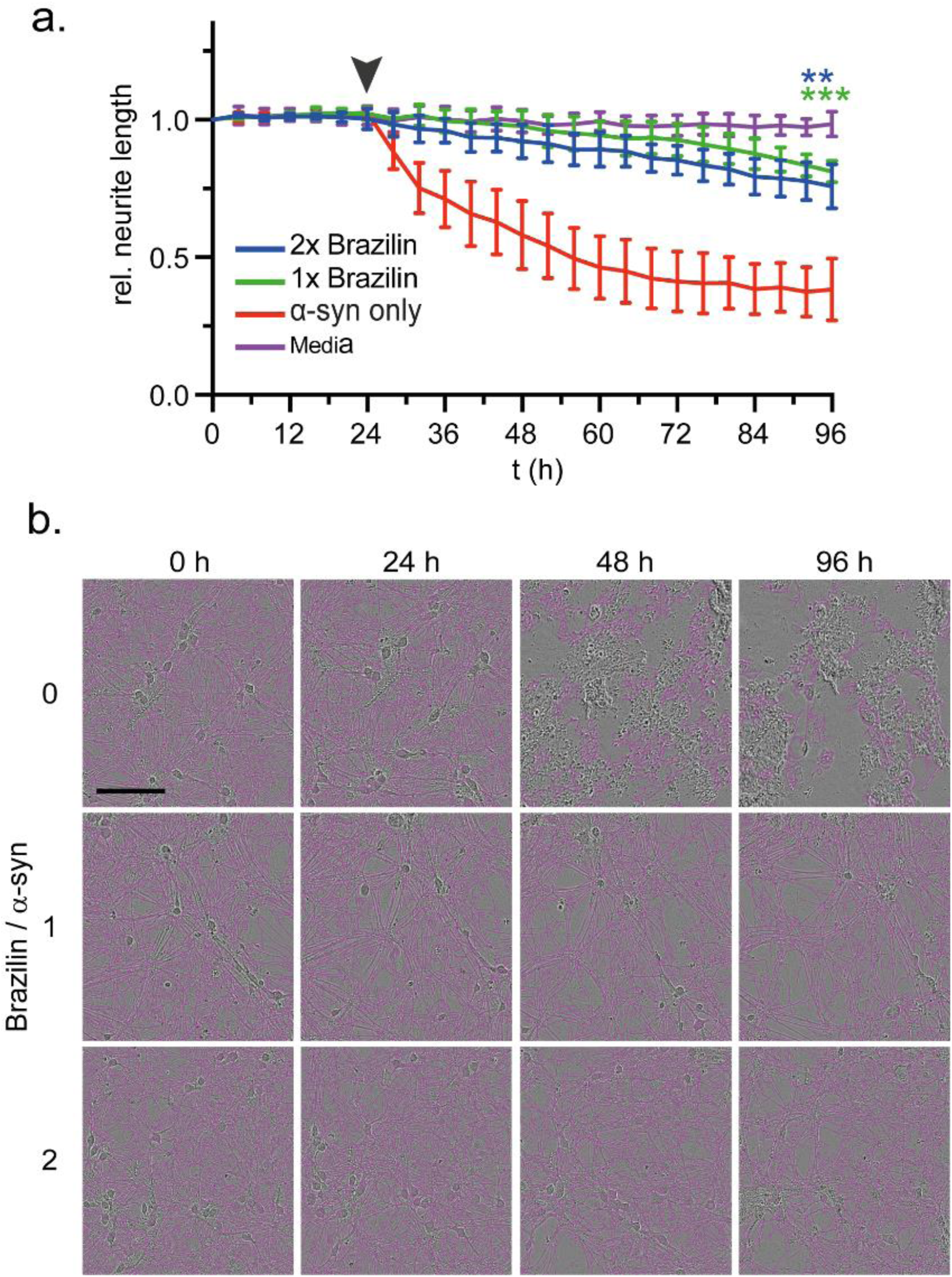
a) Neurite length of primary mouse hippocampal neurons monitored by live cell imaging after incubation with Brazilin remodeled α-syn fibrils; ** denotes p ≤ 0.01, ***denotes p ≤ 0.001 compared to untreated α-syn fibrils), means ± SD, n = 4. b) Live cell images of primary mouse neurons during incubation with Brazilin at 0x, 1x and 2x molar ratio. Pink overlays represent neurites counted. Scale bar is 100 μm.

In total, these data indicate that Brazilin treatment potently inactivates seeding-competent and neurotoxic α-syn assemblies. They furthermore suggest that, unlike polyphenols such as EGCG, but similar to Orcein and its derivatives, it does so by promoting the formation of large insoluble aggregates over smaller seeding competent species. This in contrast to Brazilin’s effect on Aβ42, where it was reported that Brazilin dissolved and remodeled amyloid fibrils into granular aggregates [47].

### Molecular Modeling of Brazilin Binding

Firstly, we explored the binding of Brazilin to α-syn monomers using molecular dynamics. The peptide structure-prediction software, RAFT [53] was used without extensive minimization to produce 1000 viable random coil structures giving an ensemble of unstructured α-syn monomers. One hundred of these were chosen at random and each simulated for 100 ns in explicit water both with and without Brazilin present, giving a total of 20 μs of trajectory data. The simulations with Brazilin were set up by docking 10 ligand molecules to the monomer chain using BUDE [55,56], corresponding to ∼13 mM ligand concentration in the simulation box. Three pairs of trajectories (with/without brazilin) were chosen at random from the 100 hundred pairs of simulations for visual examination. Final structures are illustrated in Figures 9a, 9b and Movie1. Each vertical pair of molecules in Figures 9a, and 9b started from the same initial α-syn structure. Figure S6 shows snapshots of structures taken at regular time points along one of these trajectories. The simulations demonstrate the collapse of the protein and its close association with most of the Brazilin molecules present both on the surface and interior of the polypeptide chain.

**Figure 9:**
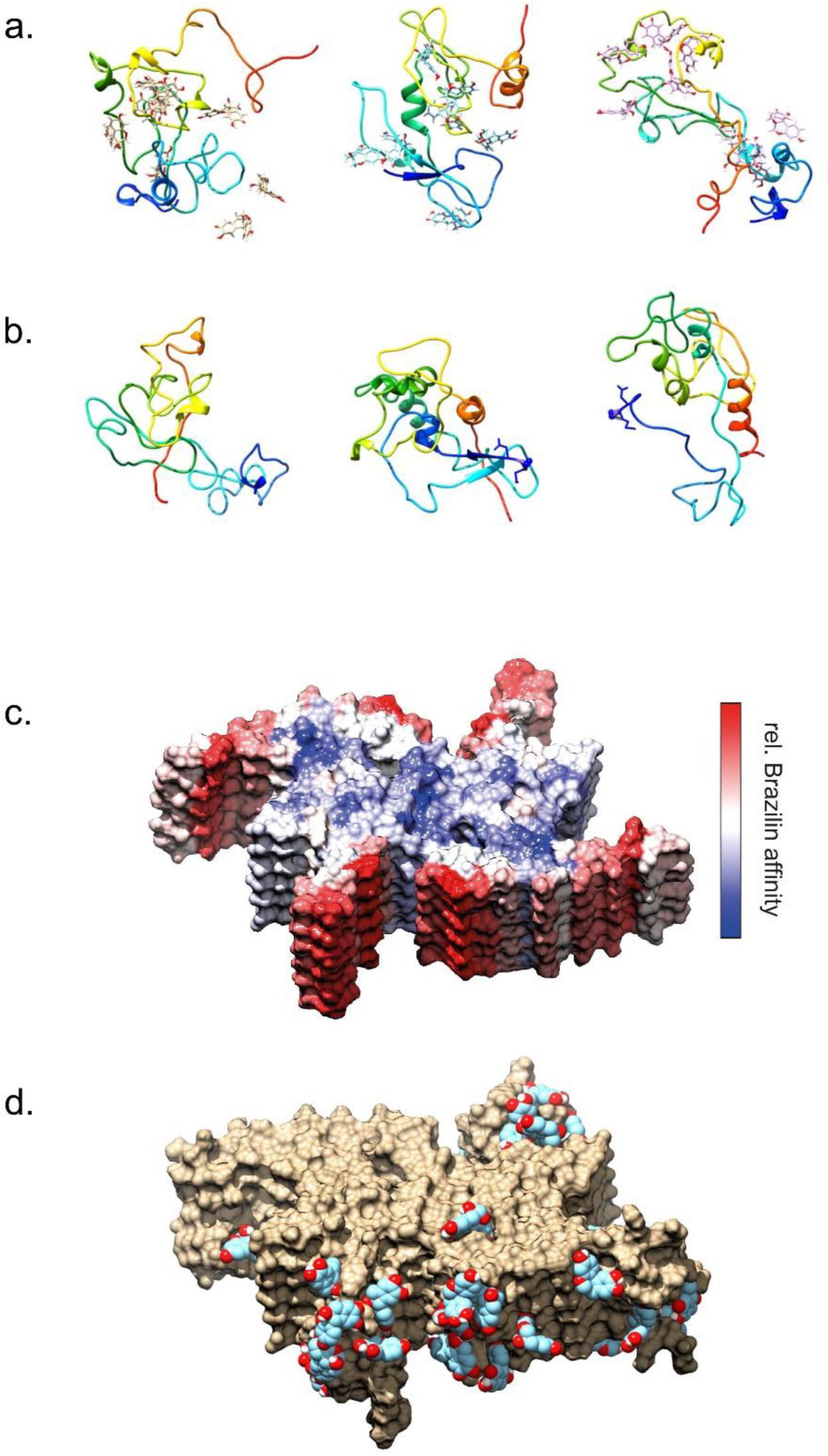
a), b) Simulated α-syn structures after 100 ns equilibration from three of the trajectories with Brazilin (a) and without (b). Each vertical pair started from the same initial structure. Protein chains are illustrated in rainbow coloring (N-term blue → C-term red). c), d) Simulated α-syn fibril fragment (PDB 6a6b) in the presence of ∼100 mM Brazilin. (c) Structure colored according to the logarithm of the number of contacts between Brazilin and the protein. Red represents maximum contacts (highest affinity) through white to blue representing minimum contacts (lowest affinity). The upper blue face (and the lower blue face, not visible) are the directions in which addition of new monomers would extend the fiber. (d) Examples of Brazilin molecules bound to highest affinity areas of the fibril model.

Quantitative analysis of the simulation data shows a rapid equilibration such that on average about 80% of Brazilin remains bound to the monomeric α-syn (Figure S7a). The presence of Brazilin has only modest effects on the evolution of the average radius of gyration of the monomer population (Figure S8) and has a slight suppressing effect on the α-helical content of the monomers (Figure S9). From these data we can deduce that Brazilin binds to the monomers without greatly influencing the structural ensemble adopted by the unfolded monomers.

Next, we looked at the influence of Brazilin on the aggregation behavior of the monomers both with and without Brazilin present. Bespoke software was used to pack 64 of the orthorhombic boxes from the starting structures of the 100 monomer simulations into a cube with the smallest volume. The resulting cubic box was 46 nm on each side and after re-soaking with model water and salt, contained nearly 10 million atoms. The size of these systems (64 α-syn monomers with and without 640 Brazilin molecules, ∼1 mM and ∼10 mM respectively) means that sampling was limited to 200 ns for both simulations. Nonetheless, these calculations produced some interesting indications of behavior. As the simulations evolve a certain amount of aggregation of α-syn is observed. Intriguingly, when Brazilin is present there is a shift towards small aggregates of two or three monomers compared with α-syn alone where more monomers are present (Figure S10). Like the individual monomer simulations, the percentage of Brazilin molecules bound to the 64 chains in this simulation is around 80%, corresponding to 8 molecules per α-syn monomer (Figure S7a, b). Movies 2a-c illustrate the initial starting structure in the box and the final structures with and without Brazilin bound. These images correspond to the aggregate-size data shown in Figure S10.

Finally, to investigate the affinity of Brazilin to a fibril structure, we simulated a fibril fragment (PDB code 6a6b) with and without Brazilin. In this case we loaded the simulation box with 222 Brazilin molecules (∼100 mM) to ensure extensive surface sampling to locate the best Brazilin binding regions on the fibril fragment. Since the α-syn monomers are truncated in this cryoEM structure, missing the first 36 and last 41 residues, these simulations are not directly comparable with the full-length monomer simulations described above. The fibril fragments are stable during the simulation as expected, with Brazilin enhancing rigidity in the structure (Figure S11). Inspection showed that Brazilin suppresses the motions of the N and C terminal regions, while the fibril core structure remains rigid in both simulations. The relative affinity of Brazilin for different parts of the structure is illustrated in Figures 9c, 9d and S12, showing that binding to the exterior of the fibril is favored over the faces where new α-syn monomers would bind to extend the fibril. Unsurprisingly, the fibril structure occludes some of the binding sites along an α-syn strand that are occupied by Brazilin in the monomer simulations (Figure S12).

## DISCUSSION

Previous studies have shown that α-syn self-assembles into fibrils via a nucleation dependent polymerization pathway [9–11]. Fibrillar α-syn is a major component of Lewy Bodies, a pathological hallmark of PD, making in vitro studies of α-syn relevant for PD pathogenesis [19,43]. Natural polyphenols are a commonly studied therapeutic intervention for Parkinson’s [40,41,45]. EGCG, which derails α-syn amyloid formation and remodels α-syn fibrils into seeding incompetent, non-amyloid aggregates, has served as a model compound for this class of molecules [34–36].

It has been shown that Brazilin can efficiently inhibit amyloid fibrillogenesis, inactivate mature fibrils and reduce cytotoxicity of both A*β* and human islet amyloid polypeptide (hIAPP), and α-syn [47,48,51]. On the surface, its mechanism seems to mirror EGCG, in that Brazilin inhibits fibril formation and is able to reduce the ThT fluorescence and toxicity of preformed amyloid fibrils [47,51,64]. However, our analysis of Brazilin’s effect on the *de novo* aggregation of α-syn, as well as its ability to remodel mature α-syn fibrils, reveals that its effect on α-syn is mechanistically distinct from EGCG and from previous reports of Brazilin on A*β* and hIAPP.

CD data demonstrate that Brazilin is able to maintain α-syn as an unfolded monomer (Figure 1). Prior research has shown that unfolded α-syn exists as a conformational ensemble around two forms, one extended and one partially compact, and that the partially compact form is a precursor to amyloid formation [30,31]. From native IM-MS data we conclude that Brazilin is binding specifically to the compact conformation of α-syn between charge states 5+-7+ (Figure 4). Interestingly, it has been shown that EGCG, and several other potent inhibitors of α-syn aggregation also bound specifically to the compact conformation, while dopamine binds specifically to the extended conformation of α-syn [21,30,65], which may suggest a common specific mechanism of polyphenolic inhibitors on α-syn monomers.

While it is important not to over-interpret the limited sampling available from the molecular dynamics simulations, both single-monomer trajectory data and simulating 64 monomers in a box yielded a binding stoichiometry around 8 Brazilin molecules per monomer that was similar to that observed in ESI-MS experiments (Figure S6). Correspondingly, Brazilin binding occurs throughout the α-syn chain, but affinity decreased near the C-terminus (Figure S12). Simulations suggest that the small molecule has little effect on the average secondary structure of the protein apart from a small reduction in alpha helicity and a corresponding small increase in coil and turn conformations (Figure S8), corresponding to experimental data from CD experiments. The observation of more dimeric and trimeric aggregates in the simulation of 64 α-syn molecules at the expense of monomers when Brazilin is present (Figure S9) is suggestive of stabilization of α-syn aggregates, again consistent with experimental data collected here (Figure 3d).

Simulation of the fibril fragment shows that Brazilin prefers to bind to the exterior surface of the fibril rather than to the two faces where recruitment of α-syn molecules would occur to extend the fibril length (Figures 9 and S12). In brief, the modelling results are consistent overall with a mechanism of amyloidosis suppression whereby Brazilin binds to α-syn monomers and small oligomers tightly enough to inhibit the formation of seeds that initiate fibrilization.

However, our data suggest that the effect of Brazilin on preformed amyloid fibrils is unlike other polyphenols like EGCG. Brazilin inactivates seeding competent assemblies, both *in vitro* and when treating assemblies in the brain of PD patients (Figures 7 and 8). While reduction in ThT fluorescence and the absence of single fibrils in AFM and EM images would suggest fibril disassembly [47,51,64], increased insoluble high-molecular weight aggregates were observed by SDS-PAGE in the pellet fraction of ultracentrifugation experiments (Figure 3). The remodeled fibrils have a similar morphology to that of untreated α-syn; but appear in larger clusters (Figure 6c), which could explain their increased SDS-resistance and their decreased seeding capacity [4]. While Liu *et al*. did not observe fibril clusters under their experimental conditions, Brazilin slightly altered the morphology preformed α-syn fibrils at 2:1 molar ratio, resulting in wider fibrils, which may be precursors of the fibril clusters we found at 10:1 molar ratio (Figure 6) [51].

Our results suggest that Brazilin inactivates seeding competent fibrils not by disassembly, but by promoting their assembly into large, inert aggregates, similar to the natural phenoxazine Orcein and derived compounds (Figure 10) [4,52]. Correspondingly, Brazilin did not directly inhibit fibril elongation when fibrils are present at large concentrations (Figure S2). Likely, formation of large aggregate clusters limits autocatalytic replication of α-syn by secondary nucleation on fibril surfaces [4]. Secondary nucleation dominates the α-syn assembly mechanism under physiological conditions [66], and which can autocatalytically replicate. These conditions are recapitulated by the RT-QuIC assay, in which Brazilin proves to be a very potent inhibitor of the replication of misfolded α-syn assemblies.

**Figure 10:**
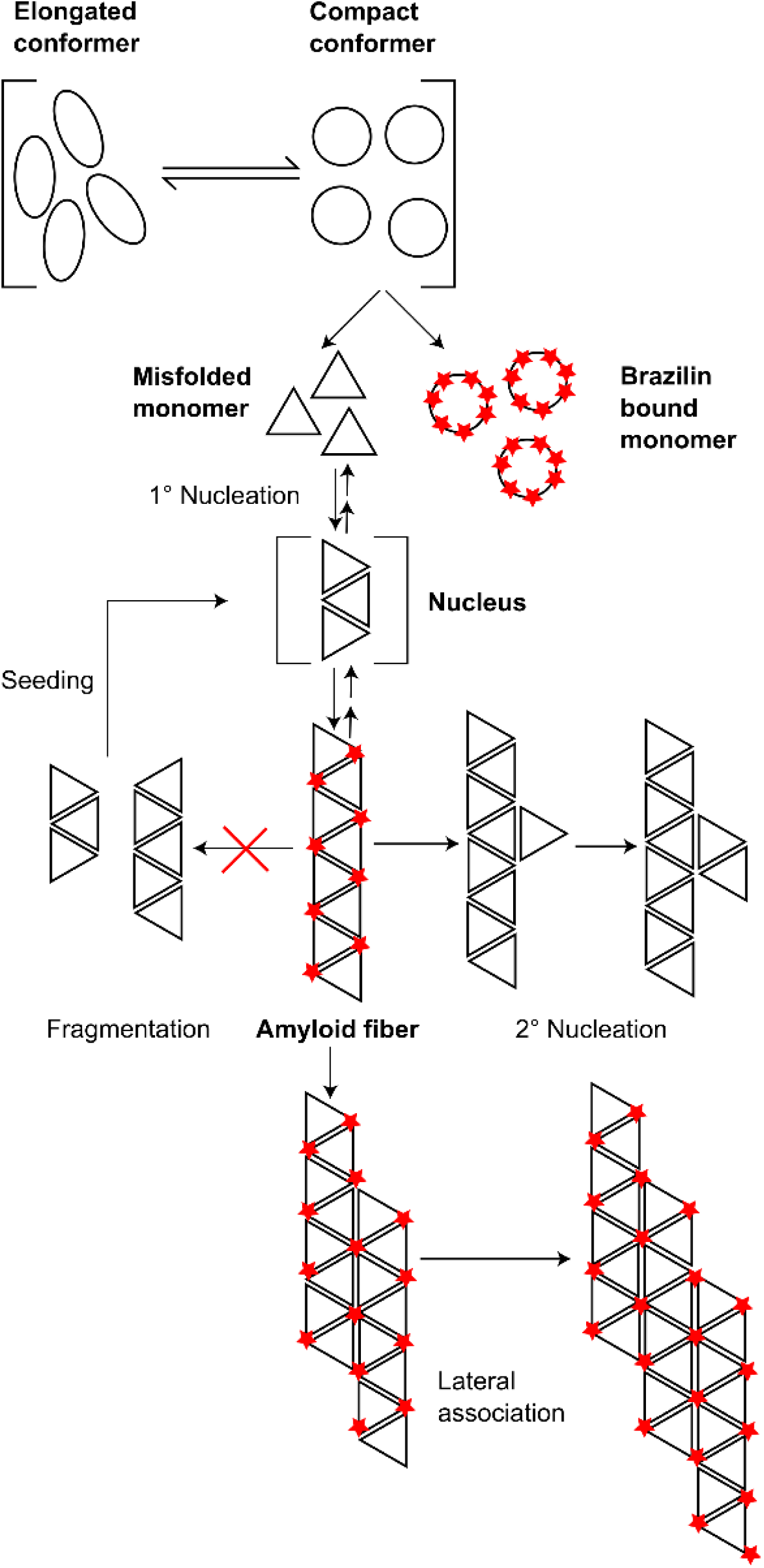
Schematic representation of proposed Brazilin inhibition mechanism.

Correspondingly, sequestering smaller amyloid species into larger aggregates removes α-syn toxicity [4,52]. This is in agreement with the reduced toxicity of α-syn fibrils after Brazilin treatment in neuronal model cells [51] which has been confirmed in primary neurons in our study (Figure 8). Our molecular modelling suggests binding of the Brazilin anions to hydrophobic pockets on the fibril surface, specifically in the vicinity of the positively charged residues K43 and K45 (Figure 9), which was also reported by Liu et al., that may promote lateral assembly of fibrils (Figure 10).

While our study cannot directly address the therapeutic potential of Brazilin, its inactivation of brain-derived α-syn assemblies suggests that it may be active *in vivo*. Previous studies on the pharmacokinetics of Brazilin found it to be highly stable *in vivo*. After a single oral dose or intravenous injection of 100 mg/kg in rats, the highest Brazilin concentrations in plasma were 82 μg/mL with half-lives of 4.5 or 6.2 h, respectively[67,68]. Brazilin has also been shown capable of crossing the blood brain barrier (BBB); after an intravenous injection of 50 mg/kg to rats Brazilin could be detected in the brain with an AUC of 340 ± 30 ng h/mL and a C_max_ of 254 ± 15 ng/mL [69]. This implies that Brazilin could cross the BBB in humans and target α-syn in a patient’s brain, making it an appropriate CNS small molecule [47]. Based on its protective activity *in vitro* and its promising pharmacokinetic properties, Brazilin may be a strong candidate as a neuroprotective and therapeutic agent in PD.

## Acknowledgements

Research was supported by the National Institute of Neurological Disorders and Stroke of the National Institutes of Health grant number 1R21NS101588-01A1 to JB and the Intramural Research Program of the NIAID- (to BC). YX, FS and SER thank the Wellcome Trust for funding (204963). Brain tissue samples were obtained from the NIH NeuroBioBank. The authors thank D. Sangar for help in protein preparation and EM imaging and S. Singamaneni (Washington University in St Louis) for use of AFM. RBS thanks the University of Bristol ACRC for access to the ARM cluster Catalyst and the GW4 Consortium for access to the ARM cluster ISAMBARD. We thank Cindi Schwarz of the NIAID Research Technologies Branch for electron microscopic imaging.

