## Supplemental Figures and Methods for "Brazilin Removes Toxic alpha-Synuclein and Seeding Competent Assemblies from Parkinson Brain by Altering Conformational Equilibrium"

**Supplementary Figures**

### Supplementary Figure S1

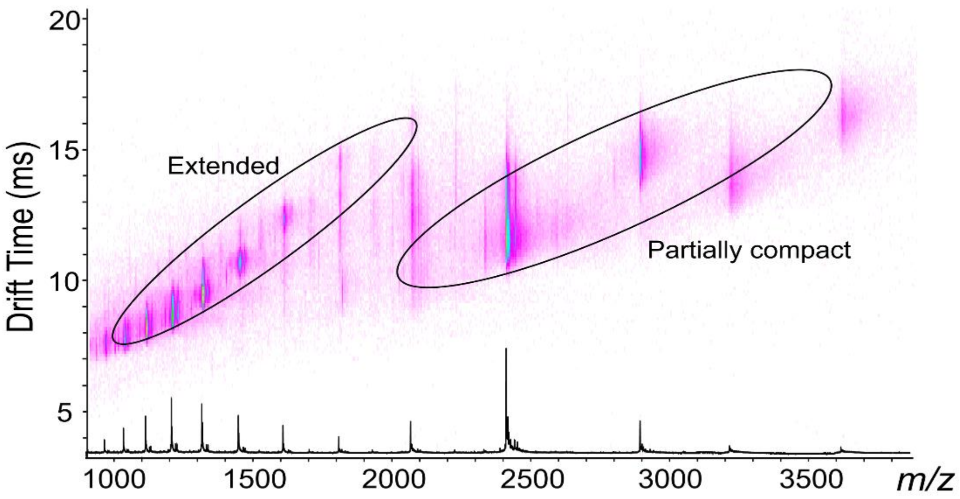


**Supplementary Figure 1:** ESI-IM-MS driftscope plot of different charge states of α-syn alone. ESI-IM-MS driftscope shows IMS drift time versus *m/z*, and the corresponding ESI mass spectrum is shown at that bottom.

### Supplementary Figure S2

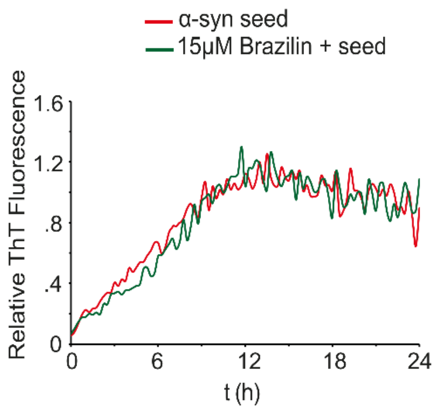


**Supplementary Figure 2:** Kinetic aggregation data of α-syn and α-syn in the presence of 15 μM Brazilin in ThT Buffer. ThT fluorescence of buffer was subtracted and ThT signals were normalized to post-aggregation amplitudes. Aggregation was seeded by 5% (w/w) sonicated α-syn fibrils. ThT fluorescence was measured in a Tecan Infinite F200 microplate reader at 436 nm excitation and 482 nm emission wavelength. Graphs represent averages of triplicate curves.

### Supplementary Figure S3

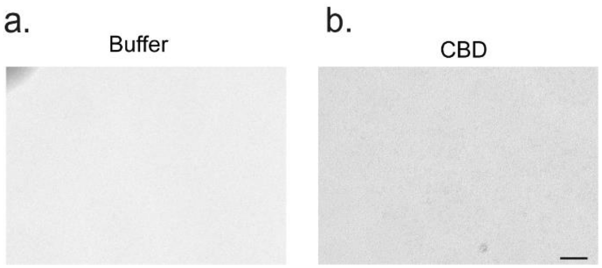


**Supplementary Figure 3:** TEM images of a) Rt-QuIC aggregation buffer (40 mM NaP, 170 mM NaCl) and b) Rt-QuIC end product when K23Q α-syn when seeded with CBD patient brain homogenate; scale bar = 200 nm.

### Supplementary Figure S4

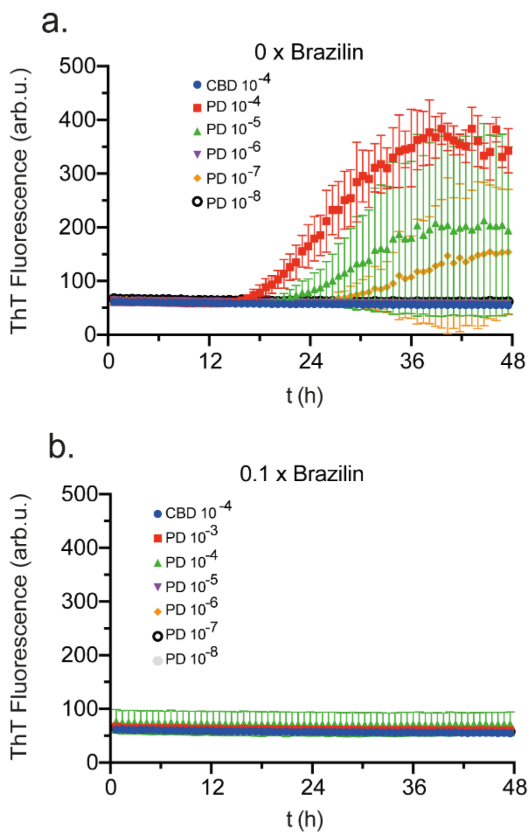


**Supplementary Figure 4:** α-syn RT-QuIC performed using K23Q α-syn as substrate, PD brain 3089 homogenate was used as seed, and CBD brain as a negative control seed. A full dilution series of brain homogenate was tested in the a) absence or b) presence of 0.1x (0.6µM) Brazilin.

### Supplementary Figure S5

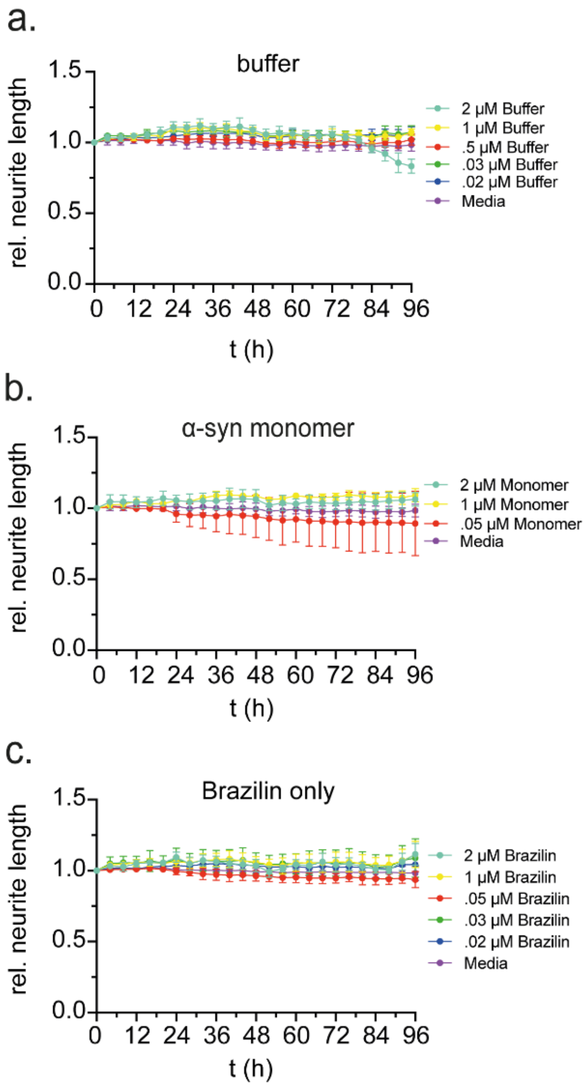


**Supplementary Figure 5:** Neurite length of primary mouse hippocampal neurons monitored by live cell imaging after incubation with a) buffer, b) varying concentrations of α-syn monomer (0.05 – 2 µM) or c) varying concentrations of Brazilin (0.02 – 2 µM). Buffer dilutions are equivalent to the α-syn concentrations 0.02 – 2 µM; means ± SD, n = 4.

### Supplementary Figure S6

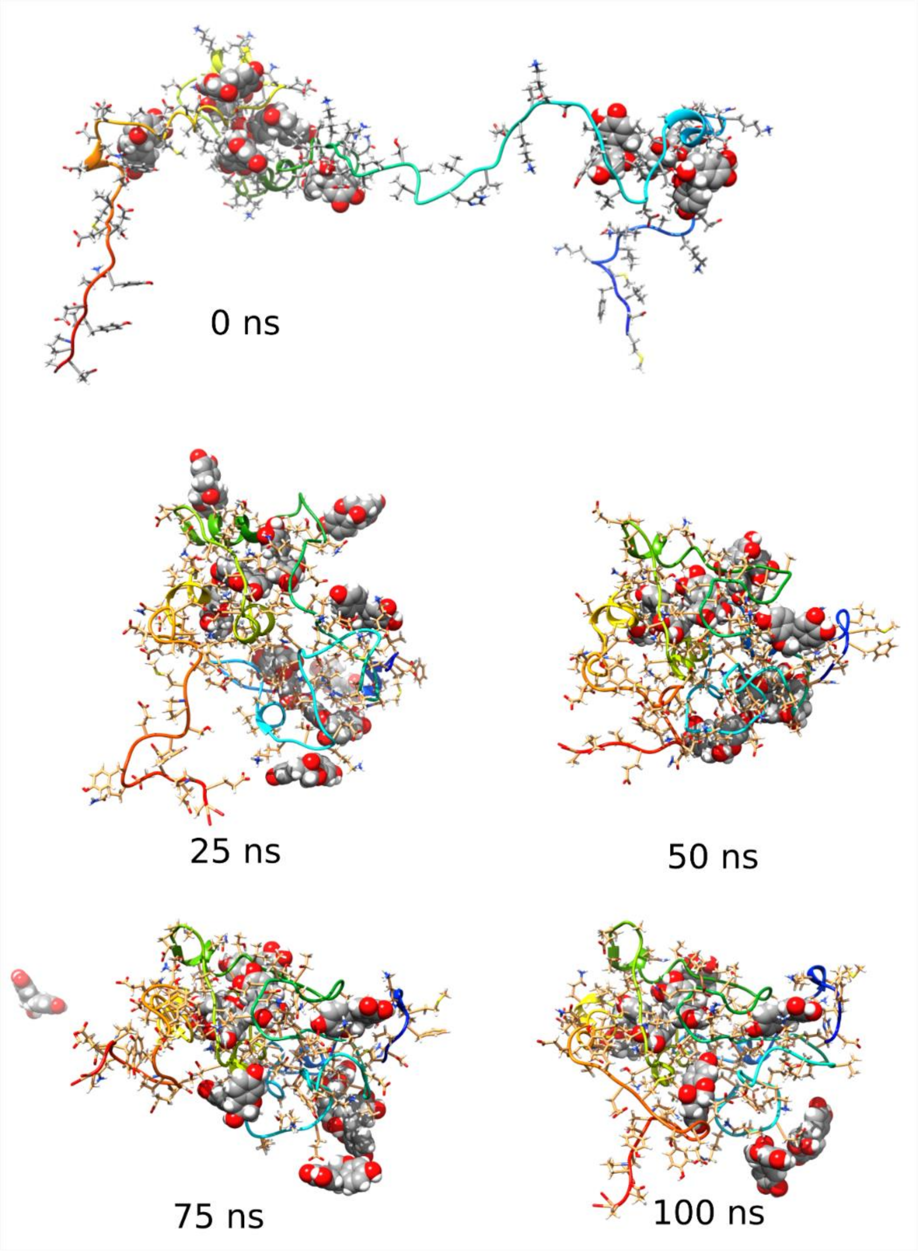


**Supplementary Figure 6:** Structures shown every 25 ns from one 100 ns monomer simulation with Brazilin. Brazilin shown as space-filling atoms, α-syn monomers shown as rainbow ribbons (blue, N-term; to red, C-term).

### Supplementary Figure S7

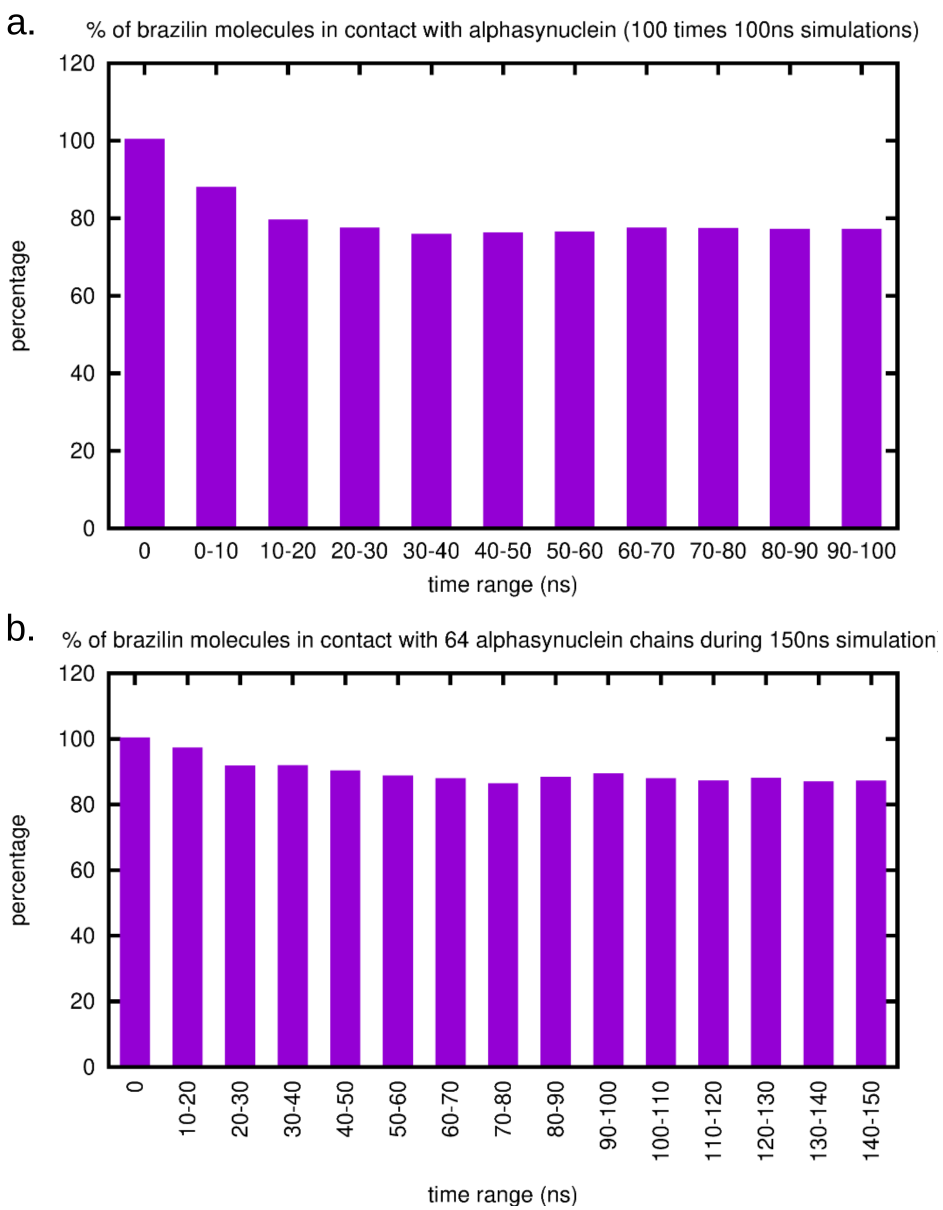


**Supplementary Figure 7:** Bar charts showing the equilibration of Brazilin binding to α-syn monomers. The simulations start from a state where each α-syn monomer is bound to 10 Brazilin molecules; a) combined data from 100 trajectories of 100 ns of a single α-syn protein; b) data from the first 150 ns of the single 200 ns trajectory of 64 α-syn monomers in a single simulation box.

### Supplementary Figure S8

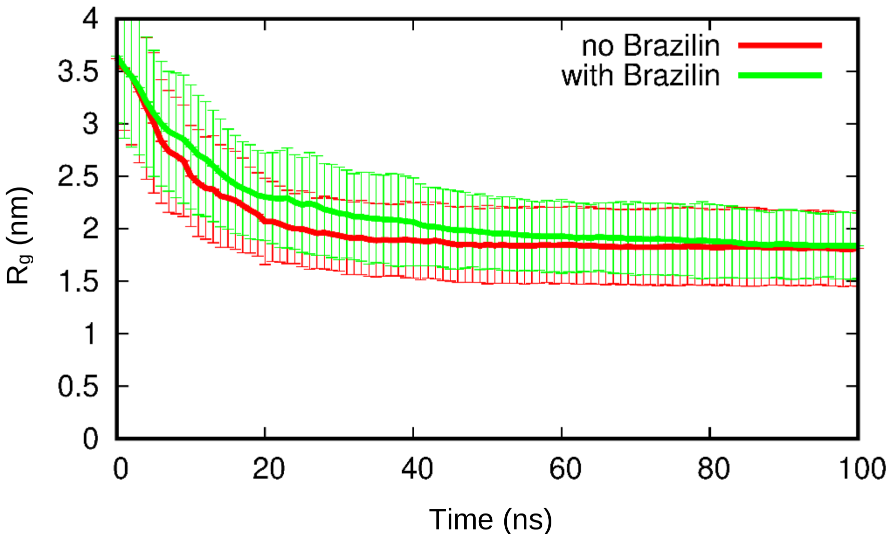


**Supplementary Figure 8:** Graph showing the evolution of the radius of gyration of α-syn monomers over the trajectories. Combined data from 100 trajectories of 100 ns of a single α-syn protein. Error bars are 1 SD from the average. Red, α-syn monomers only. Green, with 10 Brazilin molecules initially bound to each α-syn monomer.

### Supplementary Figure S9

a. α-syn

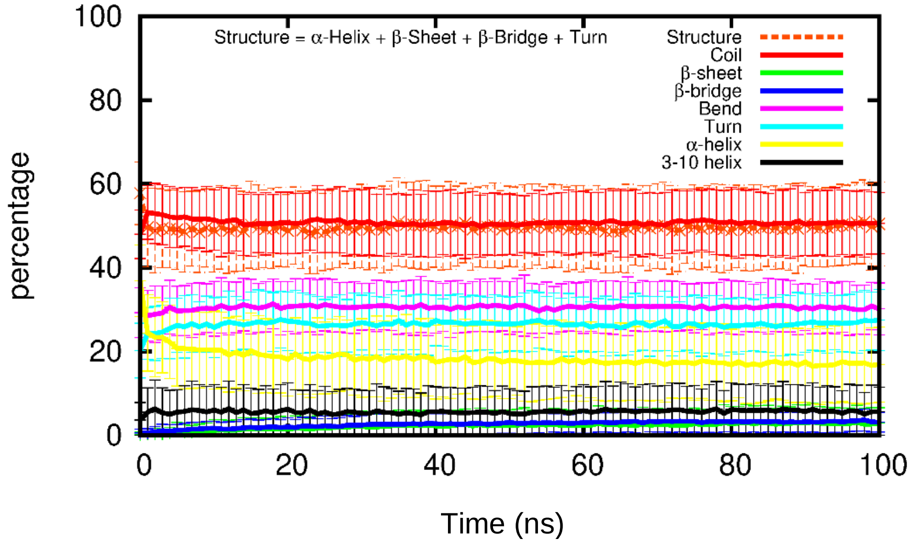


b. α-syn + Brazilin

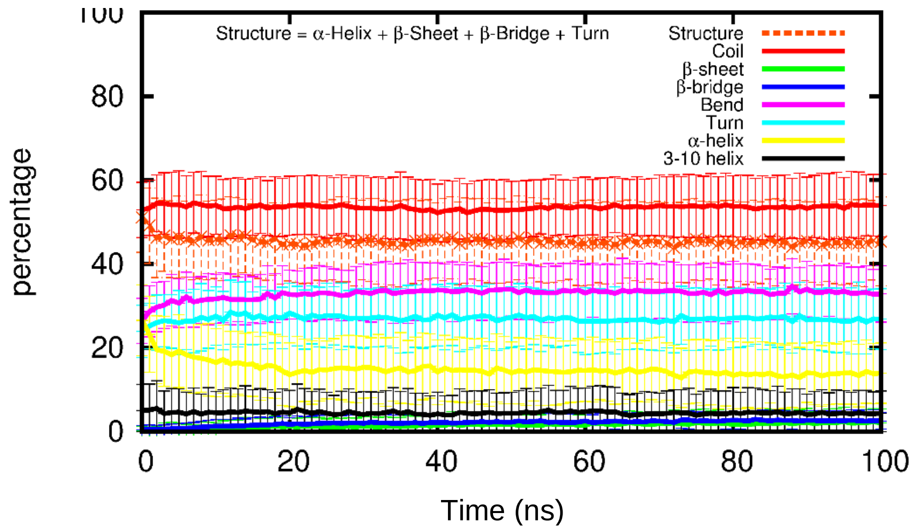


**Supplementary Figure 9:** Graphs showing the evolution of secondary structure in α-syn monomers over the trajectories. Combined data from 100 trajectories of 100 ns of a single α-syn protein. Error bars are 1 SD from the average; a) α-syn monomers only; b) with 10 Brazilin molecules initially bound to each α-syn monomer.

### Supplementary Figure S10

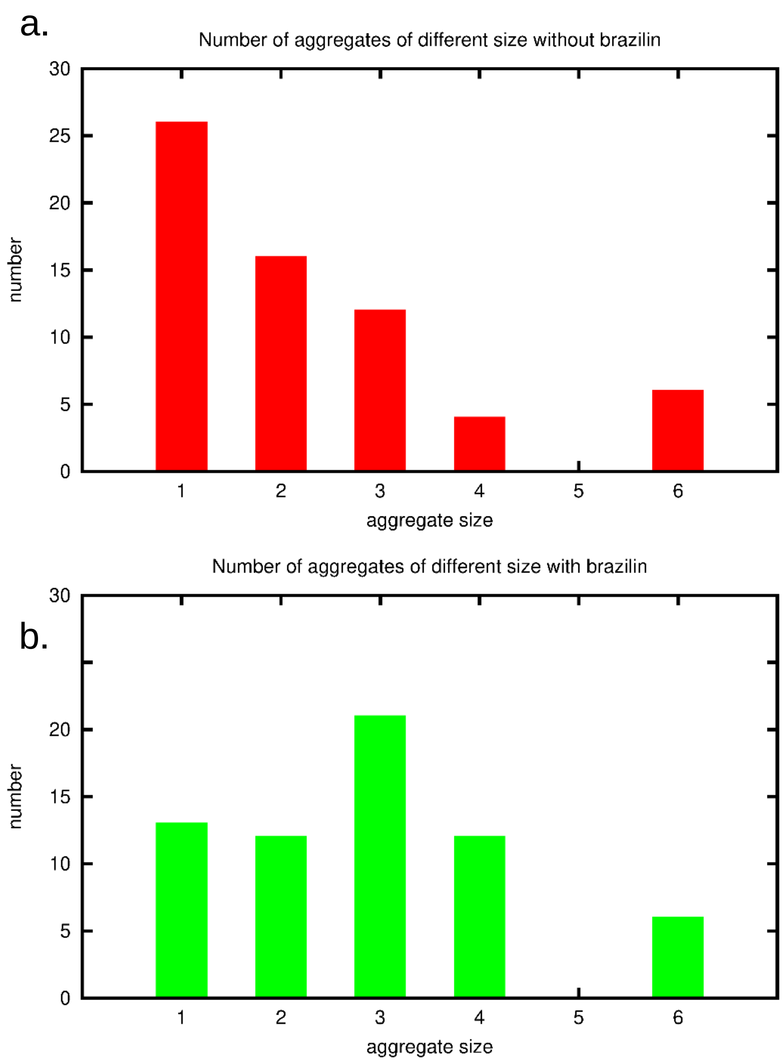


**Supplementary Figure 10:** Bar charts showing the distribution of oligomeric states of α-syn at the end (200 ns) of the simulations comprised of 64 monomers in a single simulation box. a) α-syn monomers only, b) with 10 Brazilin molecules initially bound to each α-syn monomer.

### Supplementary Figure S11

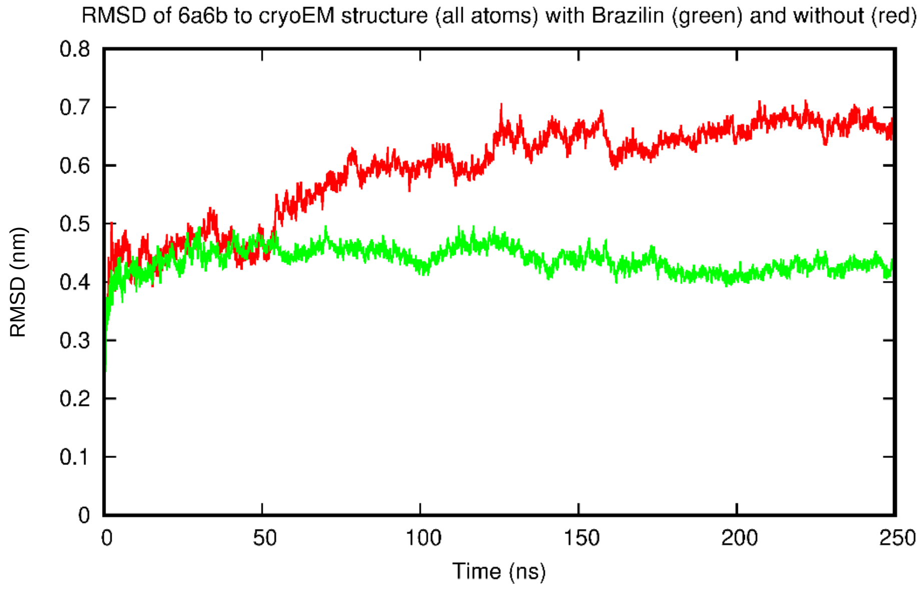


**Supplementary Figure 11:** Graph showing the root-mean squared deviation of the protein atoms with respect to the cryoEM structure (6a6b) over the simulations. Red, α-syn fibril fragment only. Green, with 222 Brazilin molecules included in the simulation box.

### Supplementary Figure S12

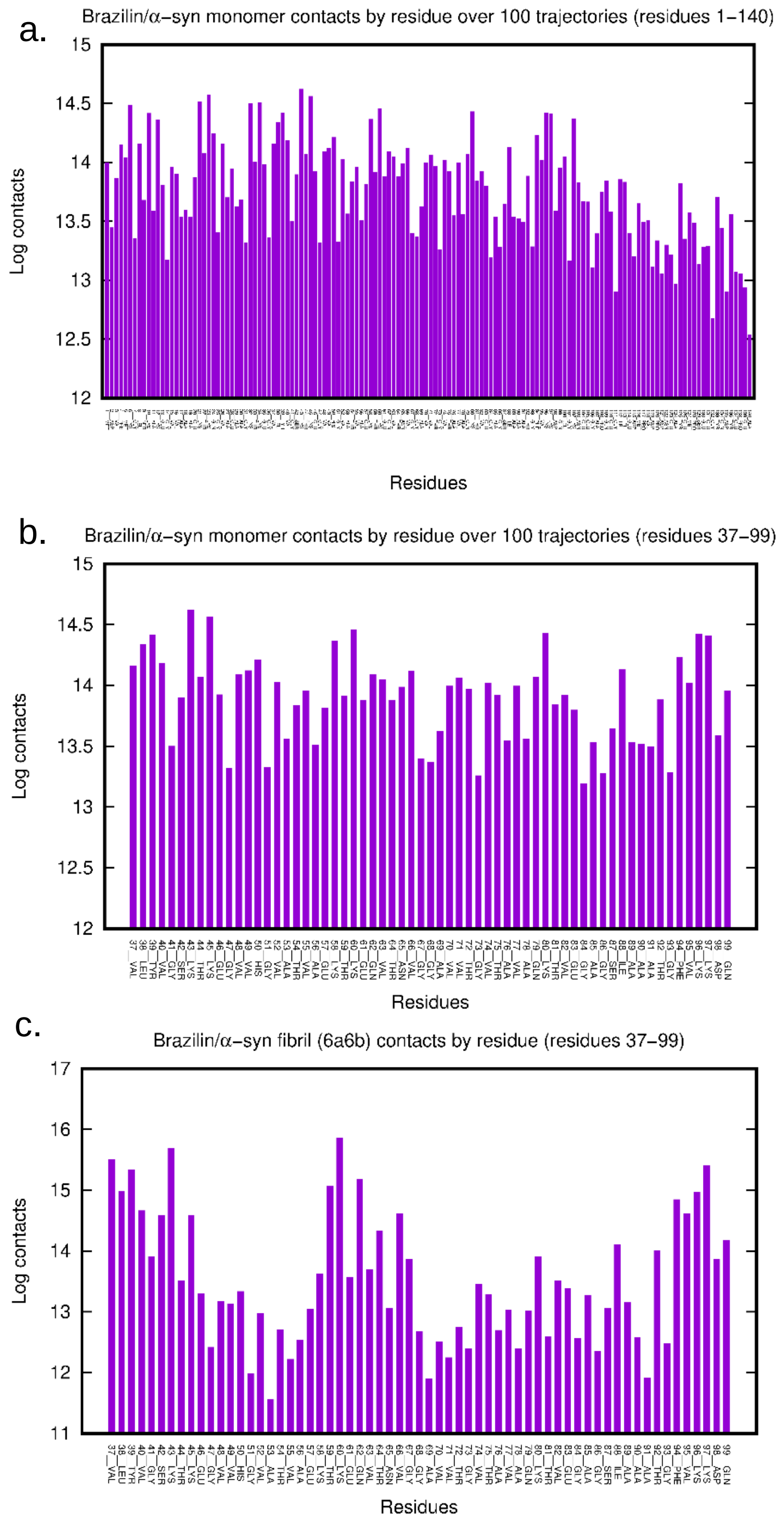

**Supplementary Figure 12:** Bar charts depicting the number of contacts made between each α-syn residue and a Brazilin molecule over the simulations. While the absolute numbers of $\log(N_{contacts})$ has no significance, each log interval approximates 1.4 kcal $mol^{-1}$ in binding energy. a) combined data from 100 trajectories of 100 ns of a single α-syn protein for the whole sequence, residues 1-140, b) expanded interval, residues 37-99, of (a) for comparison with (c), c) data from the 250 ns simulation of the fibril fragment in the presence of 222 Brazilin molecules.

**Supplementary Movies**

**Movie1:** Depicts rotating the 100 ns structure from Figure S6 and illustrates typical binding of Brazilin to the surface and interior of the collapsed α-syn monomer.

**Movie2a:** Depicts rotating the 64-monomer simulation box 360 º in the vertical axis. A) shows the initial disposition of the monomers and also depicts the case with 10 Brazilin molecules bound per monomer;

**Movie2b:** shows the final box with Brazilin present;

**Movie2c:** shows the final box when Brazilin is absent from the simulation.
